# Global Release of Translational Repression Across *Plasmodium’s* Host-to-Vector Transmission Event

**DOI:** 10.1101/2024.02.01.577866

**Authors:** Kelly T. Rios, James P. McGee, Aswathy Sebastian, Robert L. Moritz, Marina Feric, Sabrina Absalon, Kristian E. Swearingen, Scott E. Lindner

**Affiliations:** Department of Biochemistry and Molecular Biology, Pennsylvania State University, University Park, PA 16802; Huck Institutes of the Life Sciences, Pennsylvania State University, University Park, PA, 16802; Department of Pharmacology and Toxicology, Indiana University School of Medicine, Indianapolis, IN 46202; Institute for Systems Biology, Seattle, WA 98109; Huck Center for Malaria Research, Pennsylvania State University, University Park, PA, 16802; Center for Eukaryotic Gene Regulation, Pennsylvania State University, University Park, PA, 16802

**Author notes:** Corresponding Author: Scott E. Lindner, Ph.D. W230B Millennium Science Complex University Park, PA 16802 E.

**Keywords:** Translational Repression, Malaria Transmission, RNA-seq, DIA Proteomics, Proximity Proteomics

## Abstract

Malaria parasites must be able to respond quickly to changes in their environment, including during their transmission between mammalian hosts and mosquito vectors. Therefore, before transmission, female gametocytes proactively produce and translationally repress mRNAs that encode essential proteins that the zygote requires to establish a new infection. This essential regulatory control requires the orthologues of DDX6 (DOZI), LSM14a (CITH), and ALBA proteins to form a translationally repressive complex in female gametocytes that associates with many of the affected mRNAs. However, while the release of translational repression of individual mRNAs has been documented, the details of the global release of translational repression have not. Moreover, the changes in spatial arrangement and composition of the DOZI/CITH/ALBA complex that contribute to translational control are also not known.

Therefore, we have conducted the first quantitative, comparative transcriptomics and DIA-MS proteomics of *Plasmodium* parasites across the host-to-vector transmission event to document the global release of translational repression. Using female gametocytes and zygotes of *P. yoelii*, we found that nearly 200 transcripts are released for translation soon after fertilization, including those with essential functions for the zygote. However, we also observed that some transcripts remain repressed beyond this point. In addition, we have used TurboID-based proximity proteomics to interrogate the spatial and compositional changes in the DOZI/CITH/ALBA complex across this transmission event. Consistent with recent models of translational control, proteins that associate with either the 5’ or 3’ end of mRNAs are in close proximity to one another during translational repression in female gametocytes and then dissociate upon release of repression in zygotes. This observation is cross-validated for several protein colocalizations in female gametocytes via ultrastructure expansion microscopy and structured illumination microscopy. Moreover, DOZI exchanges its interaction from NOT1-G in female gametocytes to the canonical NOT1 in zygotes, providing a model for a trigger for the release of mRNAs from DOZI. Finally, unenriched phosphoproteomics revealed the modification of key translational control proteins in the zygote. Together, these data provide a model for the essential translational control mechanisms used by malaria parasites to promote their efficient transmission from their mammalian host to their mosquito vector.

## Introduction

Early embryonic development in higher eukaryotes depends on essential, maternally supplied products for early gene expression until the zygote can carry out its own gene expression program. This process, called the maternal-to-zygotic transition (MZT) (1-3), involves the proactive generation of transcripts in the oocyte ahead of fertilization that are stored in cytosolic mRNPs and are translationally repressed/silenced until fertilization occurs (4). The translation of these maternally provided transcripts is essential for early zygotic development and directly precedes zygotic genome activation (ZGA) subsequent *de novo* transcription by the zygote (3, 5).

Malaria parasites (*Plasmodium* spp.) use an MZT-like process to proactively transcribe, preserve, and translationally repress mRNAs in female gametocytes until host-to-vector transmission occurs (6-9). For this essential regulatory process, *Plasmodium* uses evolutionarily conserved RNA-binding protein regulators of translational repression, such as the orthologues of DDX6 (DOZI), LSM14 / CAR-I / Trailer Hitch (CITH), ALBA family proteins, and others (7, 10). Evidence for translational repression in female gametocytes in multiple *Plasmodium* species has come from gene-specific studies as well as comparative transcriptomic and proteomic studies, which noted strong disparities in the abundances of mRNAs and their corresponding proteins (6-14). One of the most rigorous studies, conducted by Lasonder and colleagues, presented evidence for approximately 500 putatively translationally repressed transcripts in *P. falciparum* female gametocytes (6). Based on these observations, the essential use of translational repression during parasite transmission is predicted to provide a rapid means to produce the proteins needed to adapt to the new host environment and overcome its physical barriers and immunological defenses. However, this model has not been validated at a global level for the host-to-vector transmission event, such as by examining the extent of the release of translational repression after fertilization and determining what proteins are made at very early time points.

While total comparative analyses of transcriptomic and proteomic expression of male and female gametocytes have been published in recent years, the same comparative data are lacking for zygotes (6, 12, 14). Two recent publications describe transcriptomic analyses of *Plasmodium* zygotes: a bulk RNA-seq study of *in vitro* cultured *P. berghei* zygotes, and a single-cell RNA-seq study of the early *P. falciparum* mosquito stages isolated from the mosquito blood bolus throughout the first twenty hours of infection (11, 15). These publications contribute significantly to our understanding of transcription and gene regulation by specific ApiAP2 transcription factors in this stage and highlight the control of gene expression early in mosquito stage establishment. Still, neither included a proteomic comparison to identify the proteins expressed in this stage (11, 15). To date, the only published work to define the *Plasmodium* zygote proteome was completed 15 years ago, from *in vitro* cultured zygotes from the avian malaria parasite *P. gallinaceum*, which was completed prior to the release of its genomic sequence (16). The best evidence for the release of translational repression in early mosquito-stage parasites is from single-gene studies that demonstrated that ookinete surface-expressed and secreted proteins, as well as ApiAP2 transcription factors, are critical for parasite survival and transmission (9, 11, 13, 17, 18). This key knowledge gap in our understanding of this essential host-to-vector transmission strategy could therefore be resolved with a paired transcriptome and proteome of female gametocyte and zygote stage parasites.

Translational repression in *Plasmodium* transmission stages is known to be mediated by RNA-binding proteins (reviewed in (19)). In female gametocytes, members of the RNA-binding protein DOZI/CITH/ALBA complex contribute to preserving and stabilizing specific transcripts (7, 10, 20, 21). When DOZI or CITH are absent, proper development of the zygote cannot occur. Hundreds of transcripts were identified as interactors of PbDOZI in an RNA-immunoprecipitation microarray study, and a large proportion of these bound transcripts overlap with those transcripts that decrease in abundance when *pbdozi* was deleted, indicating their abundance may be directly or indirectly dependent on the presence of DOZI (21). However, nothing is known about how the DOZI/CITH/ALBA (DCA) complex selects specific mRNAs in female gametocytes or releases them for translation in zygotes.

To address these key knowledge gaps, here we have used quantitative, comparative transcriptomic and proteomic approaches to identify translationally repressed transcripts in *P. yoelii* female gametocytes, which are then released for translation following fertilization in early zygotes. Several of these proteins are essential to establishing the new infection of the mosquito by adapting to different nutrient availability, polarizing the cell to regain motility, and initiating zygotic transcription. Notably, this is the first application of quantitative Data Independent Acquisition Mass Spectrometry (DIA-MS) proteomics to *P. yoelii*, which now provides the most comprehensive proteomes of female gametocytes and zygotes for any *Plasmodium* species. These data also reveal changes in post-translational modifications associated with female gametocytes and/or zygotes that could be used to toggle translational control across this transmission event. In addition, we have assessed how the DCA translational repression complex contributes to this regulatory control point. Using both proximity proteomics and high-resolution imaging microscopy, we have assessed potential spatial and/or compositional changes in the complex across the host-to-vector transmission point. Together, this provides the first stage-matched evidence for the widespread activation of translation of repressed transcripts in early mosquito-stage parasites and we posit that changes in the spatial arrangement and composition of DCA complex play important roles in the release of translational repression.

## Results

Collection of *P. yoelii* Female Gametocytes and Zygotes

In order to compare gene expression between female gametocytes and zygotes, we first enriched these specific life stages using flow cytometry. Female gametocytes were collected from a transgenic parasite line expressing a fluorescent tag (GFP) from a female-enriched promoter (*pylap4*), while *in vitro* cultured zygotes were selected by anti-Pys25 antibodies as this protein only becomes surface exposed following fertilization (22). The stage identity of the collected cells was validated by Giemsa-stained thin blood smears and fluorescence microscopy to ensure stage-specific enrichment (Fig S1, S2).

### Transcriptomic Profiling of Female Gametocytes and Zygotes

RNA-seq libraries were generated using four million parasites per biological replicate with the Illumina Stranded mRNA Library kit for NextSeq single-read sequencing. The resulting RNA-seq data were mapped to the PY17X reference genome (plasmodb.org, release 55) using hisat2, and differentially abundant transcripts in each stage were identified using DESeq2 with at least a 5-fold difference in abundances (Fig 1, Table S1) (23). Biological replicates are well-correlated by life stage, as demonstrated by stage-specific hierarchical clustering of sample-to-sample distance (Fig 1A). Between these two stages, 2,648 transcripts were detected in total, and nearly all transcripts (92%) detected in gametocytes were also detected in zygotes (Fig 1B), reflective of the transcripts present in the shared cytosol between female gametes and zygotes.

**Figure 1:**
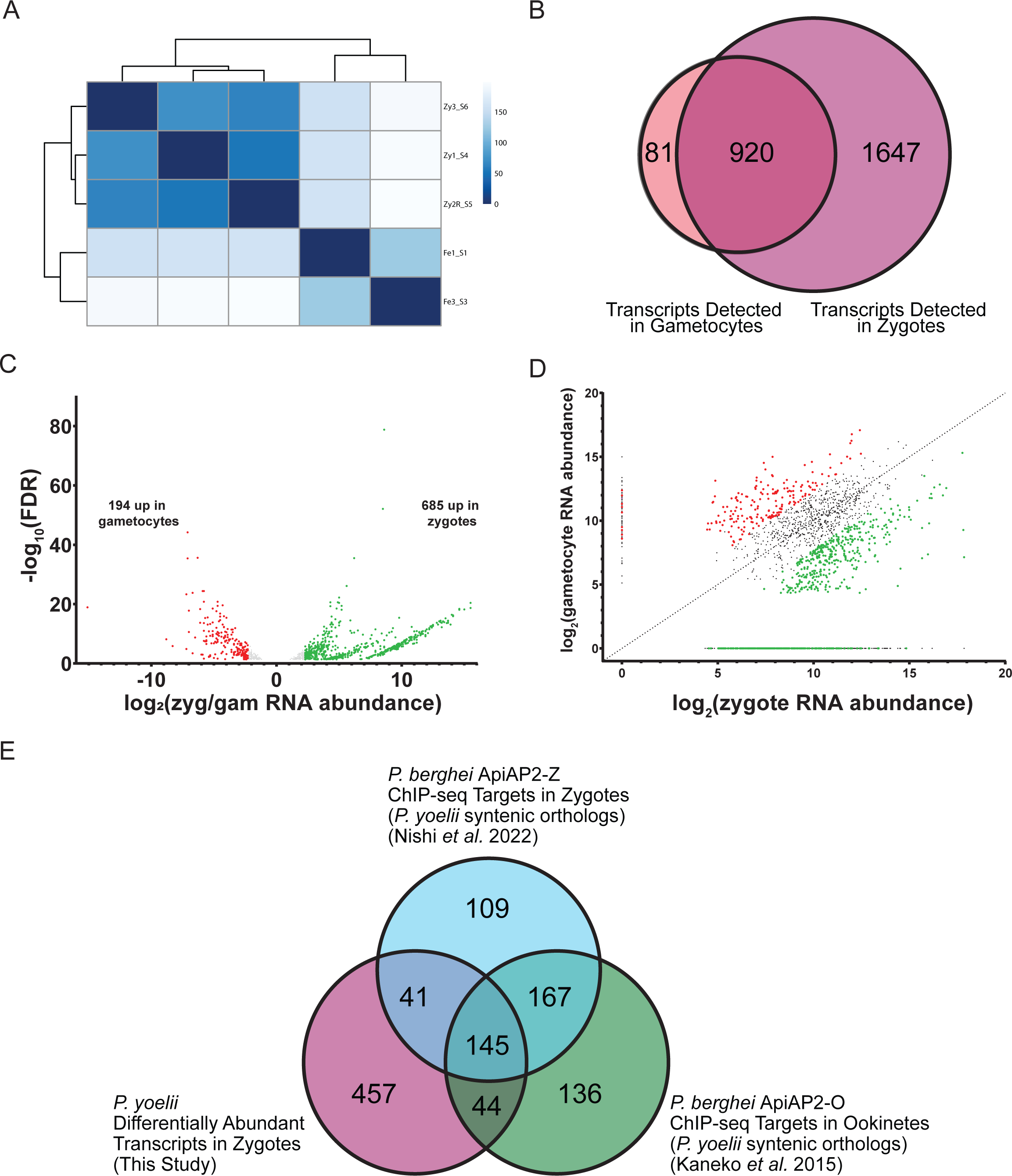
Comparative transcriptomics of *P. yoelii* female gametocytes and zygotes. A. A heatmap of sample-to-sample distance hierarchical clustering of biological replicates of RNA-seq data is shown. B. A Venn diagram illustrates the overlapping detection of transcripts in female gametocytes and zygotes. C. Volcano plot of differentially abundant transcripts enriched in female gametocytes (red) or zygotes (green). D. Dot plot of transcript abundance in female gametocytes and zygotes. Differentially abundant transcripts in female gametocytes or zygotes are highlighted as red or green points, respectively. E. Venn diagram of differentially abundant transcripts in zygotes and the *P. yoelii* orthologous transcripts that are targets of transcription factors PbApiAP2-O and PbApiAP2-Z is shown (11, 25).

Among the 194 transcripts enriched in female gametocytes include those encoding proteins involved in DNA replication licensing, like components of the origin recognition complex (ORC2), and the minichromosome maintenance complex (MCM2, MCM5, MCM6, MCM7) (Fig 1CD, Table S1). Though the female gametocyte does not replicate its DNA, the female gametocyte transcriptome was enriched in enzymes involved in DNA replication like DNA primase, DNA polymerase α and δ subunits, and components of the proliferating cell nuclear antigen DNA clamp (PCNA) and its replication factor C clamp-loader complex. It is likely that the female gametocyte is preparing for the DNA replication activities that occur early on in zygote development when the diploid zygote replicates its DNA to become tetraploid within the first hours following fertilization (10, 14, 24). Of note, many previously identified female-enriched transcripts were abundantly detected in both female gametocytes and zygotes, with no statistically significant difference in transcript abundance (*nek4*, *dozi*, *cith*, *p28*) or even a greater abundance in zygotes (*lap1*, *lap3*, *p25*).

Together, these data are consistent with the proactive transcription, storage, and stabilization of some transcripts in the female gametocyte, as well as *de novo* transcription in the early zygote that is activated by two essential transcription factors, first by the recently identified ApiAP2-Z and then by ApiAP2-O (11, 13, 25). This strategy mirrors that of the metazoan maternal-to-zygotic transition, especially as these two *Plasmodium* ApiAP2 factors are known to be translationally repressed in female gametocytes, and translated early in mosquito stage development to contribute to zygotic genome activation following fertilization (11, 13, 25). Many of the 685 transcripts that are differentially abundant in zygotes have been identified as target genes bound by *P. berghei* ApiAP2-O in ookinetes that develop from zygotes (Figure 1CD, Table S1), including the transcript for the zygote and ookinete surface protein Pbs28, one of the best studied, translationally repressed transcripts in female gametocytes (9, 25). Additionally, ChIP-seq targets of PbApiAP2-Z or PbApiAP2-O were also found to be more abundant in zygotes, including those that encode proteins involved in ookinete morphological development and secreted ookinete proteins involved in midgut traversal and ookinete-to-oocyst differentiation, such as *celtos*, *soap*, *warp*, *chitinase*, and perforin-like proteins *plp3*, *plp4*, *plp5* (Fig 1E) (11, 25). Many members of the IMC also have increased transcript abundance in zygotes, including IMC1 subunits and IMC sub-compartment protein ISP3. Together, these data support zygotic genome activation at this early timepoint and align with the reported roles of these ApiAP2 transcription factors in zygote-to-ookinete maturation.

### Establishment of Data Independent Mass Spectrometry (DIA-MS) for *Plasmodium yoelii*

Next, we applied DIA-MS proteomics in order to measure the relative protein expression between female gametocytes and zygotes. We employed a data acquisition and library-building strategy developed by Searle and colleagues (26). Unlike traditional data-dependent acquisition (DDA), in which peptide precursor ions are sampled in a serial fashion, DIA-MS fragments peptide ions over sequential wide bands of mass-to-charge ratios, thus removing DDA’s bias toward random high-abundance precursors and providing the opportunity to quantify more of the peptides present in a sample. In addition to providing deep proteome detection and robust quantification even from scarce samples, DIA-MS is well-suited to analyze obligate parasites as it is much less sensitive to interference from contaminating host proteins (26). In order to deconvolute complex spectra generated by fragmentation of wide *m/z* windows, DIA-MS data is typically searched against a spectral library, which is a compendium of MS/MS spectra for individual peptides generated from DDA experiments. In order to create the requisite comprehensive spectral libraries for our analysis beyond our empirically derived spectral data, we used Prosit (27), a deep neural network trained on millions of peptide spectra, to create *in silico* reference spectra for all theoretical tryptic peptides of the *P. yoelii* and mouse proteomes. To produce sample-specific spectral libraries tailored to the specific chromatography and MS instrumental conditions, we produced representative female gametocyte and zygote samples in sufficient quantity to enable six replicate injections of each. We then used gas-phase fractionation DIA (28), a DIA-MS technique in which narrow ranges of *m/z* are fragmented in each replicate injection, enabling deep proteome coverage without requiring offline pre-fractionation. The resulting spectra from these experiments were searched against the Prosit *in silico* spectral library and used to build empirically corrected chromatogram libraries for which the retention time and peptide fragment spectra have been empirically determined using the same experimental conditions used to generate the sample data to be searched (29). These empirically corrected spectral libraries therefore enable the quantification of more peptides than from a Prosit library alone (26).

Although the purpose of the empirically derived chromatogram libraries was to provide deeper proteomic coverage for the six gametocyte and zygote biological replicate samples with matched transcriptomic data, the libraries themselves provide valuable insight into protein expression of these parasite stages, especially since the proteome coverage enabled by gas-phase fractionation was deeper than could be obtained from the limited material available from the individual samples. A total of 3358 *P. yoelii* proteins were quantified from the combined female gametocyte and zygote library samples: 2388 from gametocyte and 3184 from zygote, with 2214 quantified in both (Fig 2A, Table S2). Each library sample represents only a single biological replicate, but by applying conservative thresholds for differential expression, we identified proteins that likely represent stage-specific protein expression. In addition to providing confirmatory evidence for the stage-specific protein expression identified from the transcriptomics-matched biological replicates, this library data set represents a valuable resource to the *Plasmodium* research community as it provides the most comprehensive *P. yoelii* gametocyte proteome to date and the first zygote proteome reported for any *Plasmodium* species.

**Figure 2:**
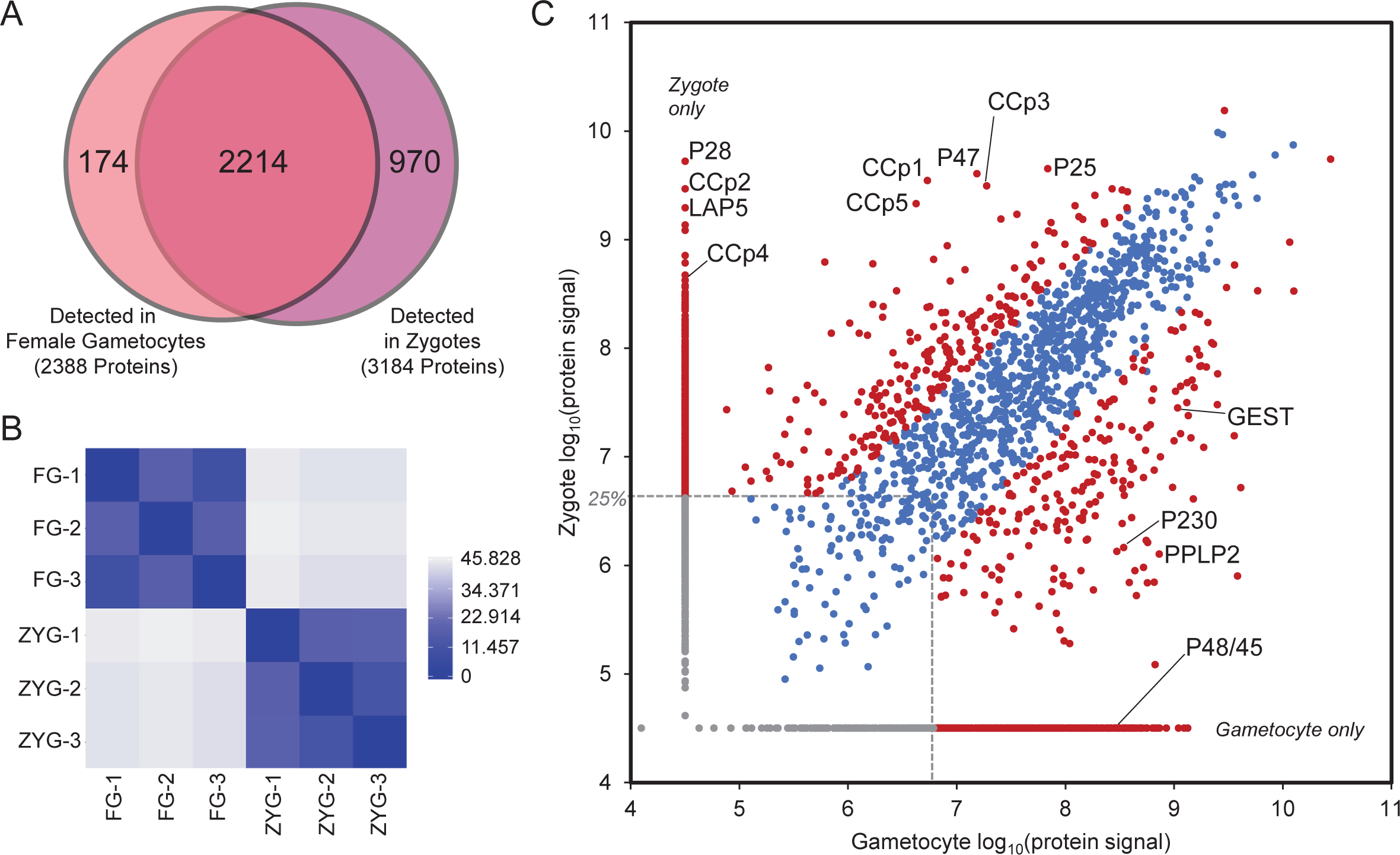
Differential protein expression between female gametocytes and zygotes. **(A)** A Venn diagram illustrates the overlapping detection of proteins from the library and biological replicate samples from DIA-MS for female gametocytes and zygotes. (B) A heatmap of sample-to-sample distance hierarchical clustering of biological replicates of DIA proteomics is shown. (C) The total signal for each protein (measured as the sum of all quantified peptide areas for the protein) from the biological replicates of samples (not the DIA-MS libraries) compared between female gametocytes and zygotes is shown. The 25^th^ abundance quartile is demarcated with dashed grey lines; only proteins detected above this threshold were considered. Red circles represent proteins that are significantly differently expressed or found in only one stage. Representative proteins-of-interest are labeled. More abundant/only detected in zygotes: gamete surface 6-cys protein P47, ookinete surface 6-cys proteins P25 and P28, and LCCL domain-containing LAP and CCp proteins. More abundant/only detected in gametocytes: gametocyte surface 6-cys proteins P230 and P48/45, perforin-like protein PPLP2, and gamete egress and sporozoite traversal protein GEST.

### Quantitative Proteomics by DIA-MS

Compiling the results of the library and samples and comparing them against the current body of published *Plasmodium* proteomes compiled in PlasmoDB (23) provides new insights into stage-specific protein expression (Table S2, Table S3). Relatively few comprehensive proteomes have been published for *P. yoelii* at any stage, so we also compared our results against published *P. falciparum* proteomes for proteins annotated as orthologs in PlasmoDB. Altogether, this compendium of *Plasmodium* proteomics includes data from one *P. yoelii* (30) (not in PlasmoDB) and 34 *P. falciparum* asexual stage samples, seven *P. falciparum* gametocyte samples, five *P. yoelii* and 20 *P. falciparum* sporozoite samples, and two *P. yoelii* liver stage samples. We now add to this list our four samples each of *P. yoelii* gametocytes and zygotes. Parsing these results reveals 276 *P. yoelii* proteins (145 of which have annotated *P. falciparum* orthologs) that were confidently identified in zygotes by at least two peptides, but for which no proteomic evidence has been detected in any other stage in *P. yoelii* or *P. falciparum*, representing a set of stage-specific protein markers for early mosquito stages. Among these are several known markers of this class, including members of the PSOP secreted ookinete proteome family (31), members of the CPW-WPC family of translationally repressed ookinete proteins (32), chitinase (CHT1, an ookinete protein essential for invasion of the mosquito midgut (33)), and the ookinete surface protein WARP (33, 34). Notably, 142 of these putative stage-specific proteins have no annotated function, representing a short list of targets for future studies to identify proteins that are essential for transmission and could thus be targetable via transmission-blocking vaccine approaches.

As with our transcriptomic data, biological replicates of the sample DIA-MS datasets clustered well by stage (Fig 2B). A comparison of protein abundance across stages identified many proteins with significantly different abundance levels or that were only detected in one of these stages (Fig 2C). A total of 550 proteins met the criteria to be considered significantly more abundant in female gametocytes (protein signal >25^th^ percentile and detected only in gametocytes, or with a ratio>5-fold and p-value < 0.05). Among these are the ApiAP2-G2 regulator of gametocyte maturation (35), proteins important for gamete egress such as PPLP2, MTRAP, SUB1 and 2, and GEST, and the gamete surface proteins (and transmission-blocking vaccine antigens) P48/45 and P230 (Fig 2C). Similarly, proteins required for licensed DNA replication that were also more abundant at the RNA level in female gametocytes were also differentially abundant. These include ORC1-5, Cdt1, CDC6, MCM2-7, MCMBP, as well as DNA primase, TOPO 1 and 2, mismatch repair enzymes, FANCJ-like helicases, RUVB1-3, and many members of the DNA polymerases alpha, delta, and epsilon, and the clamp loader complexes. In contrast, the 632 differentially expressed proteins in zygotes (same thresholds applied) include those involved in key early mosquito stage events. Among the most abundant up-regulated zygote proteins are members of the LCCL-domain containing family (CCp and LAP (36, 37)), which are essential for the maturation of gametes and ookinetes. Unlike in typical metazoans, meiosis in *Plasmodium* occurs after fertilization (38), and accordingly, we detected key meiosis proteins in the zygote, including NEK4, DMC1, and RAD51. Additionally, we detect proteins required by early mosquito-stage parasites to become motile (IMC proteins and their regulators, CelTOS), adjust their surface protein composition (p25, p28, PSOPs, CPW-WPCs, CTRP, WARP, all known members of the crystalloid), develop the ability to evade mosquito immunity (P47, Cap380), and prepare for segmentation (basal complex members). We also detected an increased abundance of NOT5, which plays a role in engaging translating ribosomes encountering non-optimal codons in the A-site (39). Moreover, we observed that key components of the mitochondria are differentially abundant, including much of the respiratory chain (Complexes III, IV, and ATP synthase), and associated transporters required for their function are differentially abundant in zygotes, reflecting the need to rapidly adapt to the lower nutrient availability of mosquitoes. In addition, transcriptional regulators ApiAP2-Z, O, and O2 are differentially abundant in accordance with their known, essential roles in early mosquito stage development. Finally, we surprisingly observed that the protein abundances for both DOZI and CITH increased in zygotes, suggesting that they may play continuing roles in mosquito stage development.

### Identification of 198 Translationally Controlled mRNAs Across the Host-to-Vector Transmission Event

Previous comparative transcriptomic and proteomic analyses of female gametocytes identified transcripts with high mRNA abundances but disparate protein abundances (6, 12). These studies did not examine translation in the subsequent zygote or ookinete stages, and so the widespread release of translational repression has not yet been demonstrated. To identify mRNAs that were translationally repressed in female gametocytes, where this repression was relieved in early zygote stage parasites, we applied several criteria. First, mRNAs must pass transcript integrity (TIN) and minimum read count thresholds as previously described (40). These mRNAs must not increase >5-fold in zygotes versus female gametocytes, as substantial increases in mRNA abundance due to the start of zygotic genome activation could explain comparable increases in protein abundance. These gates yielded 1417 mRNAs for these comparisons (Table S4). Second, proteins detected by data-independent acquisition (DIA) proteomics must either be detected only in zygotes, or must change >5-fold in zygotes versus female gametocytes. Additionally, we imposed a conservative abundance threshold (>25th percentile) to ensure that fold-change values are robust. These thresholds yielded 649 proteins for this analysis (Table S4). The union of these mRNA and protein lists identified 198 mRNAs that are translationally repressed in female gametocytes, with a subsequent release of translational control early in zygotes that permits substantial translation of the protein (black dots in Figure 3A, Table S4).

**Figure 3:**
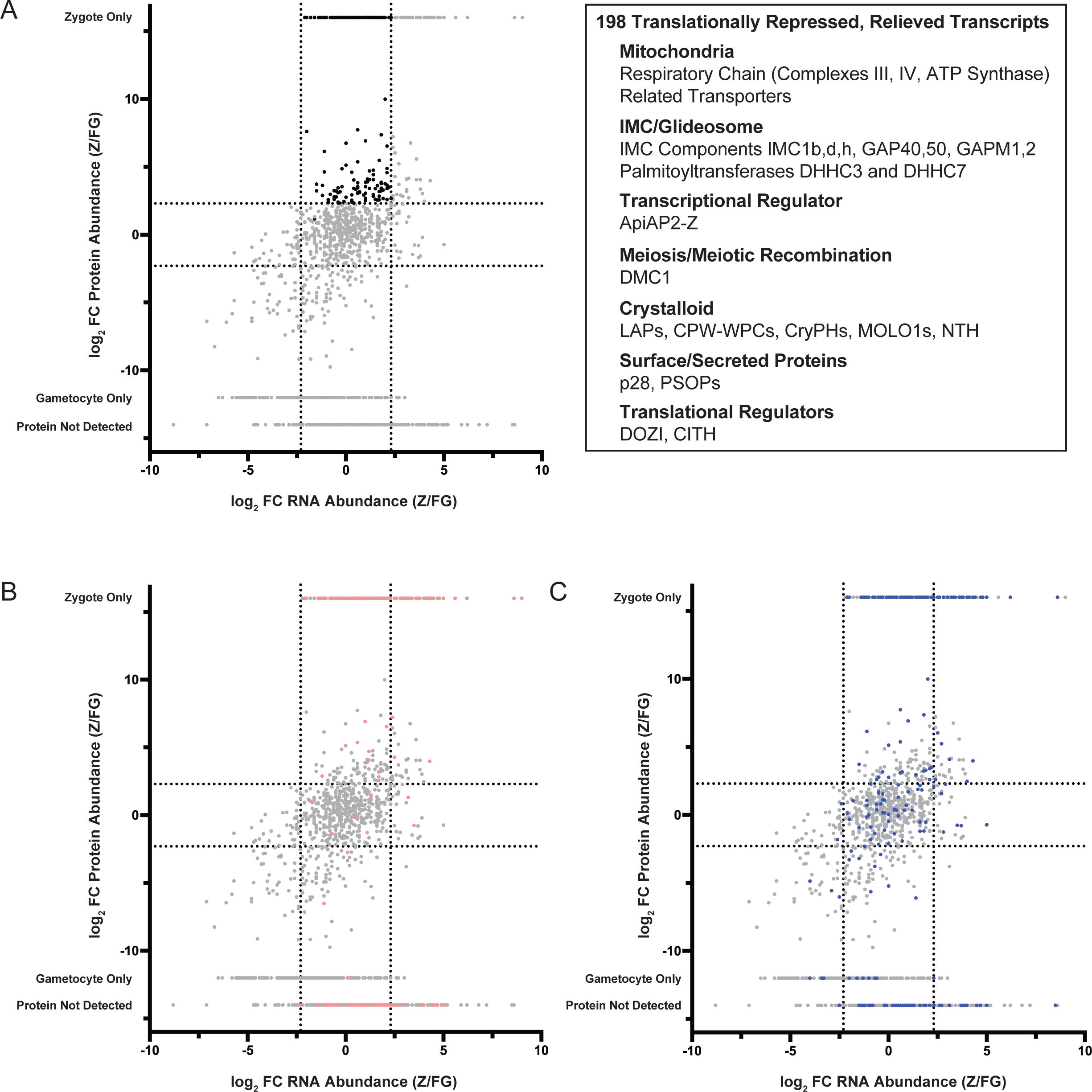
Identification of translationally repressed transcripts in *P. yoelii* female gametocytes that are translated following fertilization. **A**. The protein abundance ratio and RNA abundance ratio for transcripts and proteins detected in female gametocytes and zygotes were compared to identify transcripts detected in female gametocytes with five-fold enriched protein expression in zygotes (highlighted in black). Transcripts-of-note that are relieved from translational repression are indicated in the inset box. B. Syntenic *P. yoelii* orthologs of transcripts identified in *P. falciparum* female gametocytes as translationally repressed are highlighted in pink (Lasonder *et al.* (6)). C. Syntenic *P. yoelii* orthologs of transcripts identified to interact with PbDOZI in *P. berghei* gametocytes are highlighted in blue (Guerreiro *et al.* (21)).

Of the 198 mRNAs that are subject to translational repression in the female gametocyte that is relieved in the early zygote, there is an overrepresentation of mitochondrial proteins that function in the respiratory chain (Complexes III and IV, ATP synthase) (41), as well as related metabolic enzymes and carrier proteins. This is reflected in all three categories of GO term analyses that note statistically significant terms related to the respiratory chain, active transport across the inner mitochondrial membrane, oxidative phosphorylation, and more (Table S4). This prioritization of mitochondrial functions is consistent with the parasite’s requirement to change modes of energy production away from glycolysis during asexual blood stage growth in the glucose-rich environment of the mammalian host. While the shift toward the use of the respiratory chain begins during gametocytogenesis (42), these data indicate that specific components or overall reliance are required to a greater extent in zygotes in the more restrictive environment of the mosquito, even while still bathed in the blood meal. This finding aligns with data indicating that efficient, full respiration is required for mosquito stage development, and that atovaquone treatment of mosquitoes causes arrest of parasites at the zygote stage (43, 44).

In addition to escalating mitochondrial functions, these translationally repressed mRNAs also include those that produce proteins that are critical for core ookinete functions. These include known surface and secreted proteins of zygotes and ookinetes (p28, PSOP family proteins), regulatory proteases (plasmepsins VII and VIII), as well as many of the known crystalloid proteins (e.g., PH-like proteins, MOLO1, NTH, and CPW-WPC proteins) (45). This aligns with previous publications that identified these mRNAs as being translationally repressed in single-gene studies (reviewed in (19, 46)). Additionally, this group includes 27 members of the glideosome/inner membrane complex required for ookinete development and motility (e.g., GAPM1, GAPM2, GAP40, GAP50, IMC1b,d,h) ((47-49), and reviewed in (50)). *De novo* IMC formation is essential in order to initiate and extend a protrusion as the zygote first becomes a retort and develops further into the elongated, motile ookinete (reviewed in (51)). Importantly, two palmitoyltransferases (DHHC3 and DHHC7) known to be an IMC resident protein were also identified as being translationally repressed. DHHC3 has been shown to contribute strongly to ookinete motility, and disruption of its gene leads to a correspondingly severe decrease in the intensity of oocyst numbers (52). Moreover, DHHC7 is known to be an IMC resident of asexual blood-stage parasites, where its palmitoylation activity can be directed toward IMC members (47).

Interestingly, the mRNAs of DOZI and CITH are also translationally repressed, albeit incompletely, as their protein abundance continues to increase as zygotes are formed. While it is not known if DOZI and CITH continue to contribute a function to the parasite beyond its essential activities in the early zygote due to their complete arrest in this stage, these data would suggest that perhaps they might. Finally, the mRNA for the meiotic recombination protein DMC1 is also under this translational control. This is notable, as deletions of *pbdmc1* have no phenotype prior to early mosquito stage development, which first presents with a substantial decrease in both the number and size of the resulting oocysts (53).

In addition, only one ApiAP2 protein was produced upon the release of translational repression: ApiAP2-Z. These findings are in agreement with recent reports that defined ApiAP2-Z as being critical for zygotic functions, and is likely the initial driver of transcription during zygotic genome activation (11). In addition, several uncharacterized zinc finger proteins and RNA-binding proteins could feasibly contribute to translational control and/or zygotic genome activation. Together, these results align with the time-sensitive requirements that zygotes must meet in order to productively infect the mosquito (reviewed in (54)).

### Additional Features of Stage-Dependent Gene Expression

We also identified transcripts and proteins with increased abundances in either gametocytes or zygotes (Fig 3A). The gene products in this plot’s lower left section are abundantly expressed in female gametocytes over zygotes (Fig 3A, Table S4). This includes proteins involved in DNA licensing and replication, like DNA licensing factors MCM2-7 and MCM-BP, DNA primase subunits, replication factor C subunits, and DNA polymerases (α, δ, ε). These proteins in female gametocytes are likely generated in preparation for the early DNA replication following parasite transmission and fertilization in the mosquito midgut. In contrast, some of the most abundant and best-characterized proteins of early mosquito stage development can sufficiently rely upon *de novo* transcription following the start of zygotic genome activation (ZGA) (Fig 3A upper right quadrant, Tables S2, S3, S4). In addition, proteins that will be important later in mosquito stage development are also in this quadrant, such as the oocyst capsule protein Cap380, inner membrane complex proteins important for the structure of ookinetes as they develop (IMC1c, e, i, j, k), ookinete surface expressed proteins P47 and P25, secreted ookinete proteins WARP and perforin-like protein 4, and additional components of the mitochondrial electron transport chain (e.g., members of the cytochrome b-c1 complex, complex II subunits, and ATP synthase subunits F0, O, δ, and ε). Some of these transcripts are abundantly expressed in female gametocytes as well, but the increase in transcript abundance and protein abundance in zygotes highlights the importance of their expression ahead of ookinete development.

Finally, it is important to note that many of these abundant mRNAs do not have their encoded protein detected in either stage. This could indicate that the proteins are not detectable despite the presence of tryptic peptides that could reasonably be detected by mass spectrometry (Table S4). Alternatively, these mRNAs could remain under translational control beyond this early zygote timepoint to enable protein production at a later time in early mosquito stage development. In agreement with this possibility, one of these transcripts (*limp*) has been shown to be transcribed in starting in gametocytes and only becomes translated in ookinetes (55). We anticipate that other comparably regulated transcripts experiencing extended translational repression also are found in this dataset.

### Comparison of Translationally Repressed *P. yoelii* mRNAs to *P. falciparum* and *P. berghei* Datasets

We next compared the proteins identified in our list of translationally repressed transcripts to previously identified putatively repressed transcripts in *P. falciparum* female gametocytes (Fig 3B) (6). Many of the putatively translationally repressed transcripts identified in *P. falciparum* female gametocytes were found to be abundantly expressed as proteins in *P. yoelii* zygotes (pink datapoints in Fig 3B), and a little over a third (70) of the 198 transcripts we found to be translationally repressed were also identified as putatively repressed in *P. falciparum* gametocytes (6). This underscores the conservation of translational control over core developmental processes across *Plasmodium* species. Next, we compared our data to transcripts associated with PbDOZI in female gametocytes (21). We observed many that were abundantly detected as protein in zygotes, but also a proportion had no difference in protein abundance between gametocytes and zygotes, or even greater abundance in female gametocytes (blue datapoints in Fig 3C). This list of putatively translationally repressed transcripts of *P. falciparum* female gametocytes more closely clusters with the translationally repressed transcripts identified in our *P. yoelii* comparative study than those transcripts bound by PbDOZI. This observation may reflect the presence of non-specifically interacting RNAs in the native RNA immunoprecipitation experiment with PbDOZI that did not utilize crosslinking and highly stringent wash conditions. Moreover, this may indicate that additional transcript-specific translational controls that are independent of DOZI may be present in female gametocytes.

### Changes to the DOZI/CITH/ALBA Complex Across the Host-to-Vector Transmission Event

In order to determine mechanisms that may control translation across host-to-vector transmission, we assessed the composition of the DOZI/CITH/ALBA (DCA) translationally repressive protein complex before and after transmission. We and others have identified components of the DCA complex in *P. yoelii* and *P. berghei* via IP/MS in asexual blood stages, gametocytes, and oocyst sporozoites either with or without crosslinking (7, 10, 20). However, it was not clear how the DCA complex may respond to transmission cues that are predicted to cause the release of its bound and regulated mRNAs and thus relieve translational control.

Therefore, we employed proximity proteomics by fusing a variant of *E. coli* biotin ligase and GFPmut2 to the C-terminus of DOZI and ALBA4 as experimental bait proteins, or as an unfused control to account for highly abundant proteins in the same compartment (i.e., cytosol) (Fig S3, S4A). We chose this approach to not only determine the protein composition of the DCA complex across host-to-vector transmission, but also to determine if spatial organization of this complex changes. We selected DOZI and ALBA4 as they are known to associate with the 5’ or 3’ end of bound mRNAs, respectively (10, 20, 56), and thus provide protein proxies for the two ends of target mRNAs. We initially tested the first-generation BioID enzyme before adopting the use of the further engineered TurboID (TID) ligase that allowed for more robust labeling in a shorter labeling time, as we and others have recently shown for *P. falciparum* and *P. berghei* (Fig S4BC) (47, 57-61). To establish an appropriate duration of labeling, we assessed *in vivo* biotinylation with supplementation with 150 µM biotin for increasing time, then used total parasite lysate for affinity blots probed with streptavidin-HRP for both bait parasite lines, ALBA4::TurboID::GFP and DOZI::TurboID::GFP (Fig S4B-F). We found that one hour of biotin supplementation was sufficient for robust biotin labeling by TurboID in total parasite lysates for both female gametocytes and zygotes. Therefore, we used these labeling conditions for all experiments and analyses described here.

Biotinylated proteins were able to be captured from female gametocyte or zygote lysates on magnetic streptavidin beads for on-bead tryptic digestion and mass spectrometry analysis (Fig S4G-K). Biotinylated proteins proximal to either bait protein were captured, and their identities were inferred from peptides detected by tandem mass tag-mass spectrometry (TMT-MS) for quantitative analyses. The abundance of detected proteins was normalized to the abundance of proteins detected in cytosolic, unfused TurboID experiments to account for any non-specific biotinylation. Detected proteins greater than four-fold enriched over the unfused control were considered enriched interactors of DOZI or ALBA4 bait proteins. Three biological replicates of TurboID experiments were prepared for each bait or control protein in both gametocytes and zygotes. With these datasets, we next made a comparison of protein interactions between bait proteins and across life stages to changes to the complex pre- and post-fertilization (Table S5).

By first comparing DOZI’s or ALBA4’s interacting partners in gametocytes and zygotes, we observed that proteins proximal to either protein changed substantially across the host-to-vector transmission event (Fig 4AB). Few protein interactions remained consistent between female gametocytes and zygotes, with only 14 proteins associated with ALBA4 in both gametocytes and zygotes, while only 13 interacted with DOZI in both life stages (Fig 4CD). The 14 stage-independent interactors of ALBA4 included ALBA-family proteins ALBA1 and ALBA2, poly(A)-binding proteins PABP1 and PABP2, and RNA-binding proteins HoMu, GBP2, and DDX1, whereas the stage-independent interactions of DOZI included known interactor eIF4E and RNA-binding protein CELF2. The remaining protein interactions of DOZI and ALBA4 were stage-dependent, and perhaps these differences correspond to distinct functions in mRNA regulation in each life stage. In gametocytes, ALBA4 also interacted with proteins involved in translation at both the 5’ and 3’ end of transcripts, such as small and large ribosomal subunit proteins, initiation factors eIF4A, eIF4G, eIF3A, and RNA-binding proteins DOZI, CITH, CELF1 and CELF2, and DBP1. These associations were lost in zygotes when ALBA4 shifted to interact with proteins found at the 3’ end of mRNAs, including PABP1 (cytosolic) and PABP2 (nuclear), as well as ALBA1, ALBA2, GBP2, DDX1, DDX5, HoMu/Musashi, UPF1, and ribosomal proteins of both the large and small subunits.

**Figure 4:**
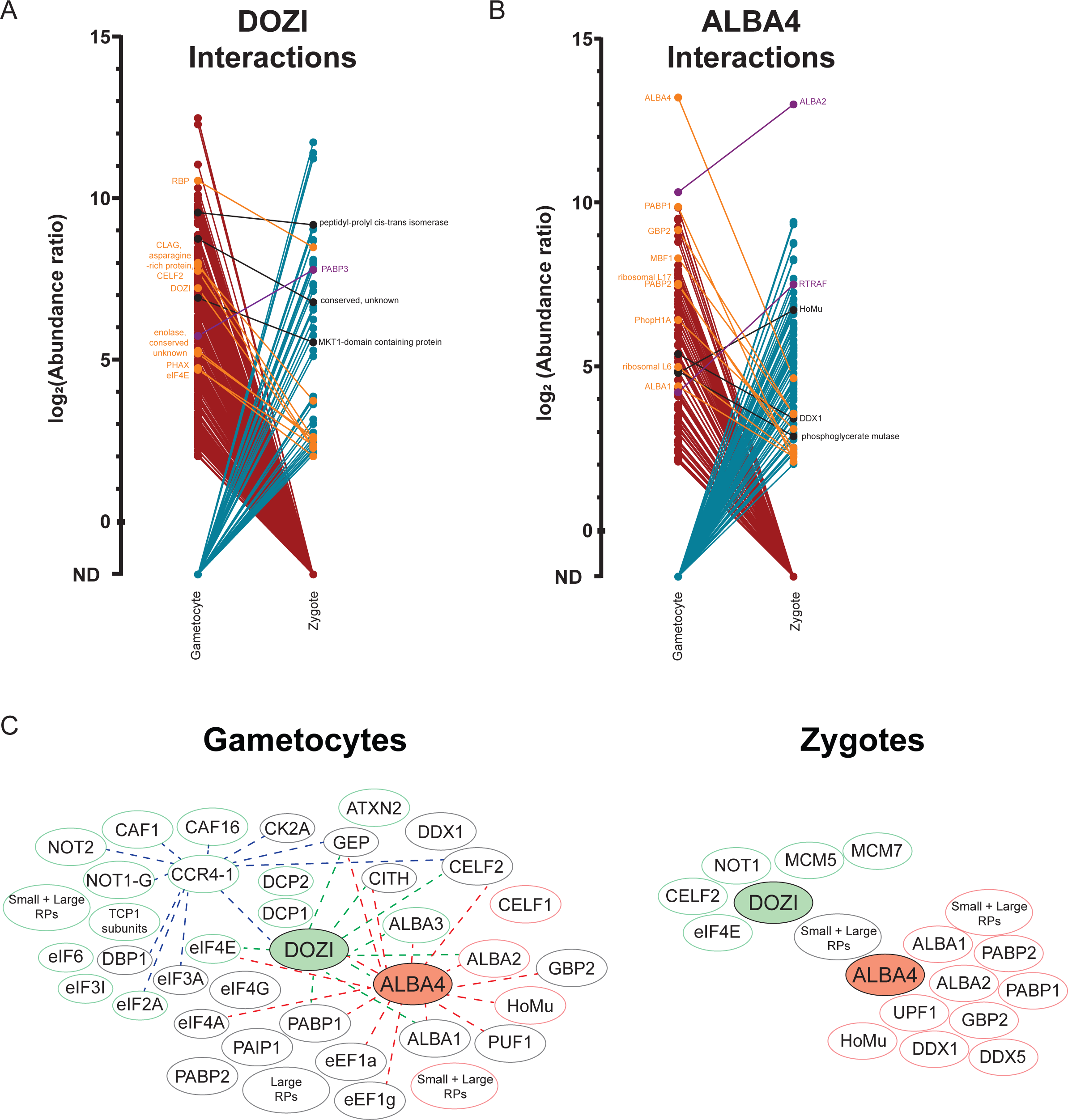
TurboID proximity proteomics reveals changes to the major translationally repressive protein complex across life stages. A and B. The abundance ratios of proteins detected over control biotinylation in gametocytes and zygotes for (A) PyDOZI::TurboID::GFP or (B) PyALBA4::TurboID::GFP are plotted. Abundance ratios were calculated for each protein as 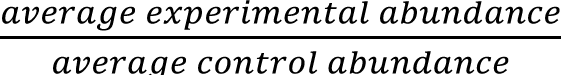. Proteins with an abundance ratio > 4 were considered enriched as interactions with PyDOZI or PyALBA4 over background labeling. C. Network of protein interactions-of-note detected in DOZI-TurboID and ALBA4-TurboID experiments in gametocytes (left) and zygotes (right). Proteins in grey circles are detected as proximal interactors of both DOZI and ALBA4, proteins in green circles are only detected as DOZI interactors, and proteins in red circles are only detected as ALBA4 interactors. Colored lines represent interactions previously detected in other immunoprecipitation-mass spectrometry experiments: green lines are interactions detected in PbDOZI IP-MS in gametocytes from Mair *et al.* (10), red lines are interactions detected in PyALBA4 IP-MS in gametocytes from Munoz *et al*. (20), and blue lines are interactions detected in PyCCR4-1 IP-MS in blood stages from Hart *et al*. (80).

We observed that DOZI also interacted with proteins found at either the 5’ and 3’ end of transcripts in female gametocytes, as well as RNA-binding proteins CITH, ALBA1 and ALBA3, factors involved in mRNA decapping (Dcp1, Dcp2), many members of the major deadenylation complex CAF1/CCR4/NOT complex (CCR4-1, CAF1, CAF16, NOT2, and NOT1-G), regulators of translation (eIF2, eIF3, eIF4E, eIF6), a homolog of ataxin-2 and the recently described FD1 RNA-binding protein (62). However, DOZI was no longer proximal to these proteins in zygotes. Most notably, DOZI exchanges its interaction with NOT1-G for one with its paralogue, the canonical NOT1 protein. Of these paralogues, only NOT1 has a bioinformatically predictable domain known to stimulate the ATPase activity and concomitant release of RNA by DOZI’s orthologues in model eukaryotes (63). These findings indicate that NOT1-G and NOT1 proteins form distinct complexes in gametocytes and zygotes, respectively, which aligns with the essential functions of NOT1-G in gametocytes and may contribute to an mRNA-release mechanism (40).

We next compared the interactions of DOZI and ALBA4 between each other to compare changes to the complex pre- and post-fertilization (Fig 4). Nearly two-thirds (62%) of the proteins proximal to ALBA4 in gametocytes were also proximal to DOZI at this stage, including many proteins known to bind to the 5’ or 3’ ends of mRNAs (Fig 4C). This indicates that DOZI/CITH/ALBA mRNPs adopt a condensed conformation during translational repression in female gametocytes. This condensed complex includes reciprocal labeling of DOZI and ALBA4, as well as mutual labeling of RNA-binding proteins (CITH, GBP2, CELF2, PABPs, ALBA1, PUF1, DDX1), large subunit ribosomal proteins, translation initiation factors (eIF3 subunits, eIF4A, eIF4G), GEP, and casein kinase 2A. Detected interactions-of-note are represented in the schematic in Fig 4CD, in which we also incorporated previously reported IP-MS data for these proteins into our protein network to indicate cross-validating data. Many previously reported interactions of DOZI or ALBA4 were also detected here by TurboID in gametocytes, including all previously detected PbDOZI core interactors identified by native IP-MS (Fig S5). In contrast, very few previously identified interactors of DOZI or ALBA4 were detected in the zygote stage by TurboID, and not many proteins were detected as interactors of both ALBA4 and DOZI at this stage (Fig 4). In fact, ALBA4 and DOZI are not reciprocally labeled in zygote stage experiments, consistent with the dissolution and/or spatial rearrangement of this complex into distinct 5’ and 3’ subcomplexes after the complex’s role in transcript stabilization and translational repression in gametocytes is completed.

Taken together, the compositional and spatial rearrangements detected by proximity proteomics are consistent with DOZI/CITH/ALBA mRNPs 1) adopting a condensed state during translational repression in female gametocytes, and then 2) undergoing composition changes that promote its elongation in zygotes following the release from translational repression. These data align with recent smFISH and KARR-seq studies of mRNA compaction in model eukaryotes that reached similar conclusions (64-67).

### Validation of Protein Colocalization by Ultrastructure Expansion Microscopy and Structured Illumination Super-Resolution Microscopy

To further investigate subcellular protein interactions detected in the condensed DCA storage granules present in female gametocytes, we assessed the colocalization of protein interaction pairs by ultrastructure expansion microscopy as it now routinely enables sub-100nm resolution (U-ExM, Fig 5) (68, 69). We enriched PyDOZI::GFP female gametocytes as above and stained expanded parasites with antibodies against GFP and either PyCITH, PyPABP1, or PyALBA4. We observed extensive colocalization of DOZI with all three proteins in classic cytosolic puncta (pink = colocalization) (Fig 5) (20, 70, 71). Furthermore, to enumerate the colocalization of each antibody pair, we used structured illumination super-resolution microscopy (SIM, Fig S6) to determine their overlap coefficient (0 being no overlap and 1 being complete colocalization) (72) (Fig S6). As with U-ExM, we observed substantial colocalization between DOZI/GFP, ALBA4, and PABP1. In addition, because we detected PyNOT1-G as being proximal to DOZI in female gametocytes, we assessed its colocalization by SIM and also found substantial overlap in its localization with DOZI, ALBA4, and PABP1. Together, these results corroborate the proximity proteomics results and support the presence of a condensed, cytosolic DOZI/CITH/ALBA granule interacting with NOT1-G in female gametocytes.

**Figure 5:**
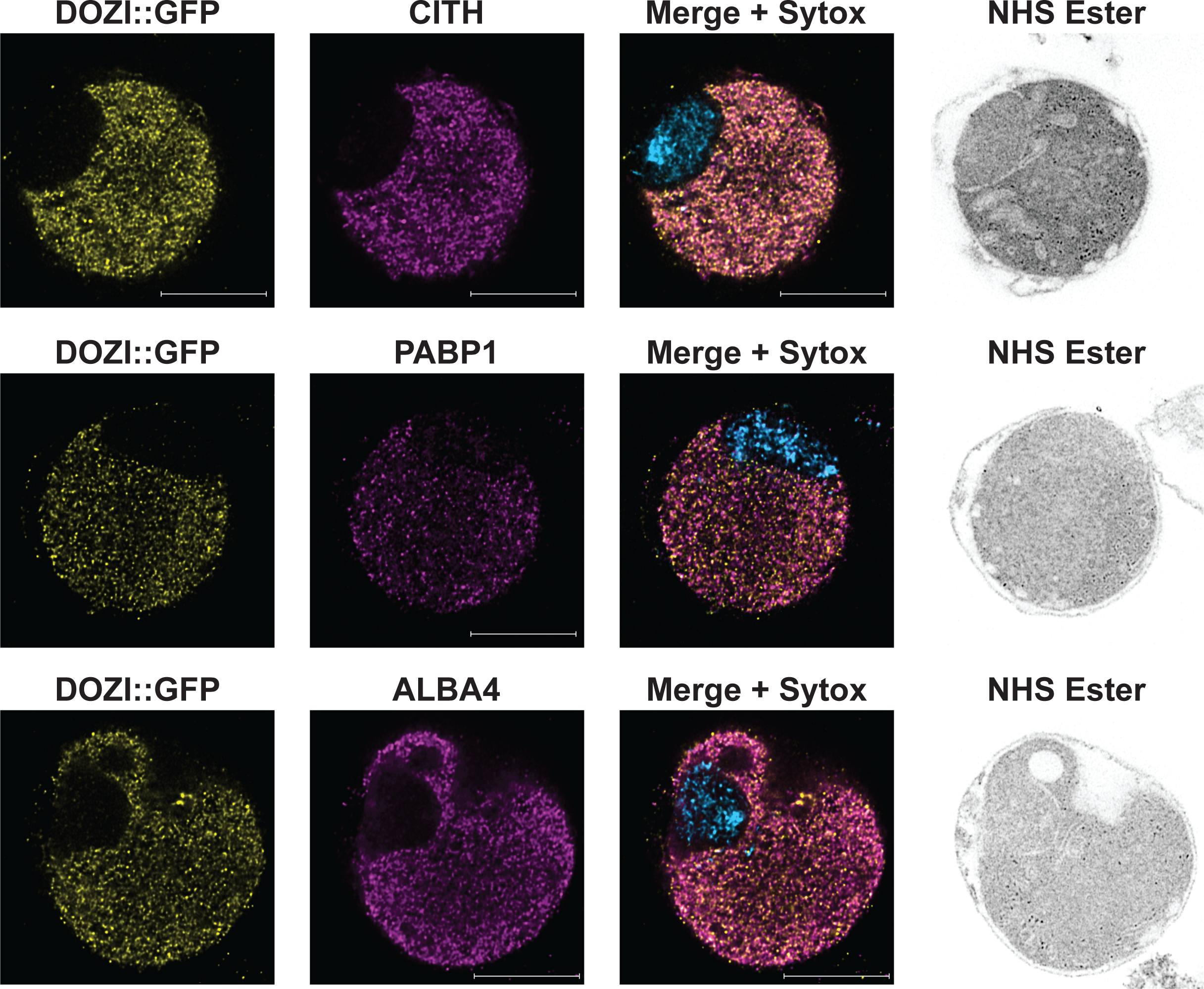
Ultrastructure expansion microscopy cross-validates protein-protein interactions in female gametocytes. PyDOZI::GFP-expressing female gametocytes were visualized via ultrastructure expansion microscopy (U-ExM) as previously described (68, 99). Parasites were stained with anti-GFP to mark DOZI (yellow) and were counterstained with custom rabbit polyclonal antibodies raised against PyCITH, PyPABP1, and PyALBA4 (magenta). Nucleic acids and proteins were stained with Sytox Far Red (blue) and NHS-Ester (gray), respectively. Colocalization is denoted with pink coloration in the merged image of a single z plane. Scale bars = 2.4 microns.

### Notable Post Translational Modifications Detected in Female Gametocytes and Zygotes

Consistent with prior reports, we have identified a wealth of kinases that are expressed in female gametocytes and zygotes (73). Some of these kinases have previously been shown to functionally regulate the essential activities of these stages (74). The quantitative proteomics data we generated in this work did not measure post-translational modifications (PTMs) because the program used to generate the *in silico* spectral libraries used for searching the DIA data is not capable of predicting spectra for peptides bearing PTMs. In order to find evidence of phosphorylation in the proteomic data, we used the DISCO tool from the Trans-Proteomic Pipeline (75) to transform the DIA data into pseudo-DDA data that could be searched for variable modifications using standard shotgun proteomics data analysis tools. Even without the enrichment of phosphopeptides, we qualitatively identified 464 phosphopeptides from 232 proteins (Table S6). Notably, all phosphorylation events on detected glideosome/IMC proteins are only seen in zygotes, with the lone exception being ISP1, which has phosphopeptides only detected in female gametocytes (Table S6). Similarly, phosphorylation of DOZI/CITH/ALBA complex members was skewed to be substantially or only detected in zygotes (Table S6). Of the 177 phosphopeptides detected in proteins associated with DOZI and/or ALBA4 in female gametocytes and/or zygotes, two-thirds (117/177) are only detected in zygotes. These phosphopeptides are detected in DOZI, CITH, CELF2/Bruno, ALBA proteins 1-4, GBP2, PREBP, DBP1/DDX3, eIF3a, eIF4a, PABP1, PAIP1, NOT1-G, CAF16, and GEP. Another 15% of the total (27/177) were observed in both female gametocytes and zygotes. These qualitative data are consistent with a phosphorylation-based regulation of the members of this complex, such as on the positionally conserved Ser20 of DOZI within a predicted intrinsically disordered region in its N-terminus (76). While these findings coincide with the release of its translationally repressive activity, establishing whether and which modifications are causal remains to be determined.

## Discussion

Malaria parasites must detect and respond to environmental cues across their developmental cycle, and perhaps most pressingly, must do so during their two transmission events between host and vector. Failing to quickly respond reduces or eliminates the parasite’s ability to persist and perpetuate itself post-transmission.

Therefore, *Plasmodium* parasites proactively prepare mRNAs ahead of time so that their encoded proteins can be rapidly produced when needed without the time required for *de novo* transcription to occur. Moreover, this provides an energetically favorable approach to preparation, as the energetic cost of translation is deferred. Together, this just-in-time approach helps the parasite to rapidly initiate the next phases of its development and countermeasures against its host or vector. This scenario mirrors the requirements of a newly fertilized metazoan zygote, which must also rapidly initiate development upon fertilization even before zygotic genome activation (ZGA) and widespread *de novo* transcription occurs. To achieve this developmental need, metazoans rely on the process of maternal-to-zygotic (MZT) transition whereby mRNAs are translationally repressed in the female gamete and then translated upon fertilization. In both *Plasmodium* and metazoans, translationally repression of select mRNA is enacted by well-conserved RNA-binding proteins (RBPs) and their partners that localize these mRNAs into cytosolic storage granules.

Much has been done to identify the RBPs that govern translational repression in *Plasmodium* parasites (reviewed in (19)). These proteins, such as DOZI, CITH, ALBA family proteins, PUF proteins, and more, constitute a weak point in these essential regulatory functions that can be exploited. In addition, several studies have identified mRNAs that putatively are translationally repressed in female gametocytes or sporozoites. In one recent study, we have also experimentally validated the translational repression and then release of specific mRNAs globally in both *P. falciparum* and *P. yoelii* sporozoites (77). These findings revealed that multiple orthogonal programs of translational control can be imposed simultaneously and relieved independently. However, such a global approach had not yet been taken for host-to-vector transmission to assess the release of translational repression during the development of female gametocytes to fertilized zygotes.

Therefore, in this study, we have used a multi-omic comparative transcriptomics and the first application of highly sensitive quantitative DIA-MS proteomics to *Plasmodium yoelii* to interrogate the control of gene expression in female gametocytes and zygotes. Several key observations now allow us to understand these stages more clearly. First, our proteomic study of zygotes provides an important and comprehensive proteomic view of this stage. Prior to this work, the only proteome of zygotes was studied by Vinetz and colleagues in 2008 using *P. gallinaceum* without a reference genome for this species (16). Our proteome greatly expands what was detected (3184 vs. 966 proteins) and fills in key details of zygote biology. Second, these data reveal what proteins can be made ahead of transmission in the female gametocyte, and also indicate which proteins cannot. For example, female gametocytes proactively prepare both transcripts and proteins required for licensed DNA replication, although they would not be required until after fertilization. Third, these data indicate what RNAs and proteins can be produced solely after transmission, without the need for the benefits of translational repression of mRNAs. These include several essential ookinete proteins (SOAP, WARP, etc.) that are rapidly transcribed and translated in the early zygote. Many of these differentially abundant mRNAs in zygotes are putative targets of the ApiAP2-Z and ApiAP2-O specific transcription factors, which are subject to translational control presumably to enable rapid *de novo* transcription of these genes (11, 25). Fourth, many abundant transcripts are detected in both female gametocytes and zygotes that were not detected as a protein in either timepoint in our study. This may reflect further repression that is not relieved until later in mosquito stage development (e.g., ookinetes, oocysts). Moreover, DIA-MS enabled the detection of post-translational modifications without the need for enrichment of these modifications. We observed several differential phosphorylation marks on members of the DOZI/CITH/ALBA complex between female gametocytes and zygotes. In other eukaryotes, such PTMs can promote regulatory events by affecting homotypic oligomerization and non-uniform scaffolding interactions that are used to form and dissolve cytosolic granules (78).

Finally, these data reveal 198 mRNAs that are translationally repressed in female gametocytes that are then translated soon after fertilization in zygotes. This enables the rapid production of proteins that are integral to essential processes of the early mosquito-stage parasite, which include the mitochondrial respiratory chain (Complexes III, IV, V) to presumably adapt to the more nutrient-poor conditions of the mosquito, and the inner membrane complex (IMC) required for proper development and polarization of the zygote, retort, and ookinete. Moreover, the translational repression program also enables rapid translation of ApiAP2-Z to promote zygotic genome activation, DMC1 for meiosis, and several well-characterized surface proteins important for ookinete viability and function. Surprisingly, we also found that DOZI and CITH have sustained mRNA expression and increased protein expression in zygotes, suggesting that they may play roles in mosquito stage development, which aligns with the timing of their deletion phenotypes (7, 10).

Our data also validate the presumed translational control of mRNAs observed in previous studies (6, 74). In one of these studies, Sebastian and colleagues identified CDPK1 as a regulator of the release of translational repression. Of the 65 transcripts that they identified to be controlled by both CDPK1 and the DOZI/CITH complex, only 9 are present in the 198 regulated mRNAs identified here. This is in agreement with validation experiments conducted by this group, which identified that not all DOZI/CITH-regulated transcripts were governed by CDPK1. This would suggest that other release mechanisms are present and could substantially contribute to TR release in the early zygote stage. Moreover, we identified mRNAs for which there is no associated protein detected in either female gametocytes or zygotes. One of these mRNAs, *limp*, has been shown to undergo extended translational repression that lasts all the way to the ookinete stage (55). Our dataset indicates that this form of extended translational control is more widespread instead of *limp* being an atypical outlying example. This model is consistent with our previous work that demonstrated the multiple orthogonal programs of translational repression can be imposed and released at different developmental times in sporozoites/early liver-stage parasites (77). Determining why the parasite would enact this complex, energetically costly gene regulatory approach is intriguing, as at face value it is counterintuitive to do so if *de novo* transcription could produce sufficient mRNAs at the time they are needed.

In addition, we sought to determine mechanisms by which the essential regulatory DOZI/CITH/ALBA complex responds to cues that promote the release of translational repression of mRNA to which it binds. Using TurboID-based proximity proteomics of DOZI and ALBA4, we found that substantial spatial and compositional changes occur across the host-to-vector transmission event. Our findings align with recent reports using smFISH or KARR-seq that refute the classic closed-loop model of mRNA translation and instead show that the translation of mRNAs correlates with a loss of their condensation in both native and stressed states (64-67). Similarly, our proximity proteomic data are consistent with the model of a condensed mRNP under translational repression in female gametocytes with both 5’ and 3’ ends of the mRNA being in close proximity, which converts to an elongated state with 5’ and 3’ ends being spatially distinct during translation in zygotes. Advanced imaging microscopy approaches using both ultrastructure expansion microscopy (U-ExM) and structured illumination microscopy (SIM) provided cross-validation of these proximity proteomics results, which showed extensive colocalization of both 5’ and 3’ end-binding proteins in female gametocytes. Finally, these data also uncovered a compositional change that this complex undergoes as the parasite develops from a female gametocyte to a zygote. We have recently shown that *Plasmodium* uses the NOT1-G paralogue of NOT1 for essential functions of both male and female gametocytes (40). Here, we found that there is a handoff between these paralogues that could provide a mechanism for translational control and its release that correlates with the timing of the *pynot1-g^-^* phenotype.

Therefore, we propose a model encompassing these multi-faceted approaches used by *Plasmodium yoelii* to promote its efficient host-to-vector transmission (Fig 6). *Plasmodium* selectively represses the translation of 198 mRNAs until after an early point post-transmission when they are necessary, but also permits the production of other proteins ahead of time that presumably are tolerated or not detrimental to the parasite in female gametocytes. The environmental stimulus/stimuli that trigger the release of translational repression is not yet known but results in a substantial reorganization of the DOZI/CITH/ALBA complex from a condensed mRNP to an elongated form when translation occurs. While this explains much of the translational control occurring during the transmission of female gametocytes, multiple lines of evidence indicate that DOZI-independent regulation is also at play. This could allow for multiple tiers of translational release over time, translation of specific mRNAs in distinct sub-localizations to promote co-translational interactions and more. Finally, we hypothesize that these same principles of translational control will apply to other *Plasmodium* species, given the substantial overlap in regulation observed between *P. yoelii* and *P. falciparum* female gametocytes.

**Figure 6:**
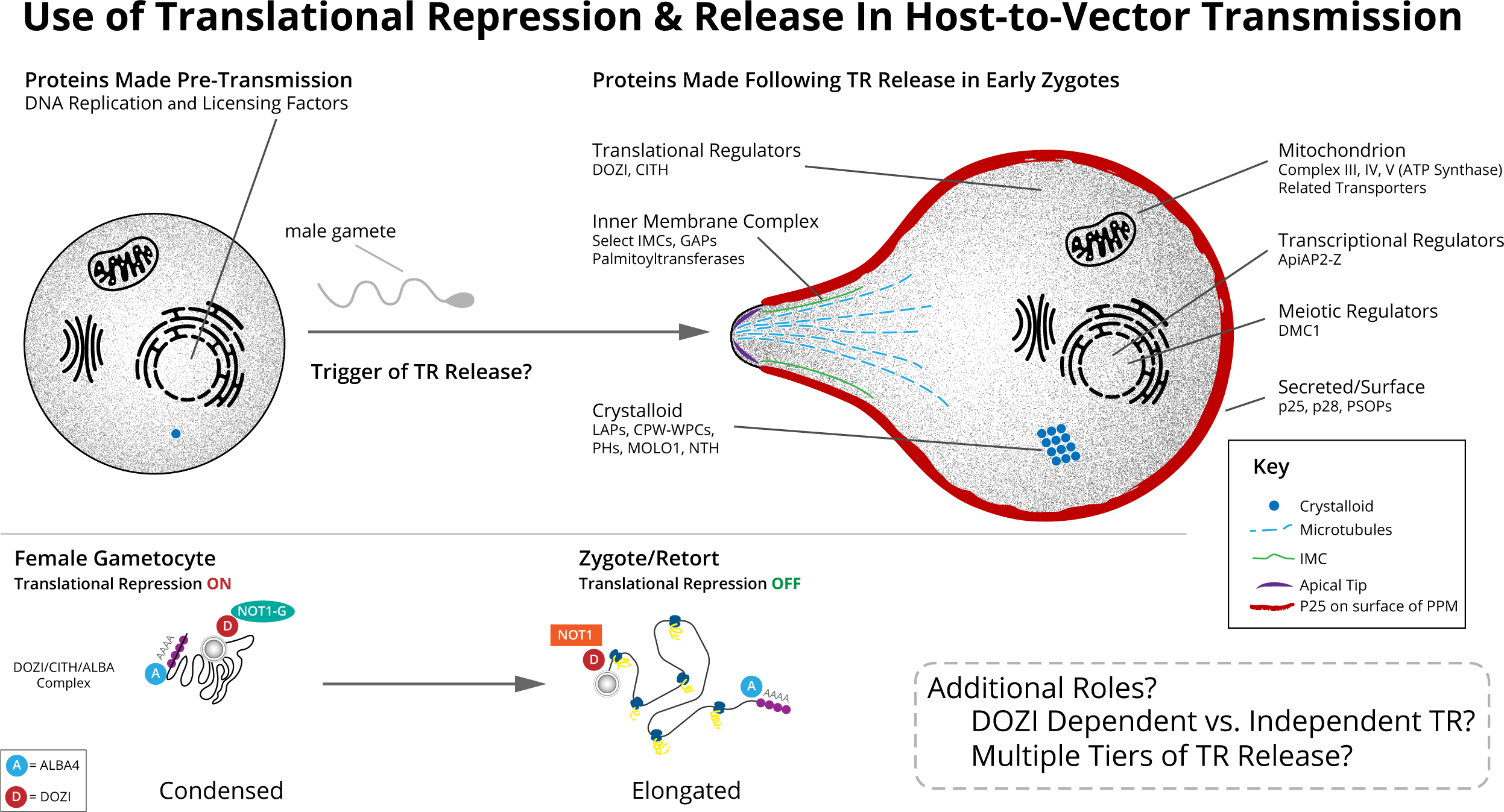
A model of the regulation of translational repression across the host-to-vector transmission event. (Top) Proteins that can be made ahead of transmission in female gametocytes (left) or that are made shortly after fertilization and formation of a zygote (right) are illustrated. (Bottom) Proximity proteomics of DOZI and ALBA4 that associate with the 5’ and 3’ end of mRNAs, respectively, revealed conformational changes in bound mRNAs between female gametocytes and zygotes, consistent with recent reports that demonstrated this by smFISH in other eukaryotes. Widespread compositional changes in the DOZI/CITH/ALBA complex are also evident, especially the handoff of DOZI interactions from NOT1-G to NOT1 across the transmission event.

## Materials and Methods

### Ethics Statement

All vertebrate animal care followed the Association for Assessment and Accreditation of Laboratory Animal Care (AAALAC) guidelines and was approved by the Pennsylvania State University Institutional Animal Care and Use Committee (IACUC# PRAMS201342678). All procedures involving vertebrate animals were conducted in strict accordance with the recommendations in the Guide for Care and Use of Laboratory Animals of the National Institutes of Health with approved Office for Laboratory Animal Welfare (OLAW) assurance.

### Use and Maintenance of Experimental Animals

Six- to eight-week-old Swiss Webster female mice from Envigo were used for all experiments performed with *Plasmodium yoelii* 17XNL strain parasites and were housed in groups of up to 5 animals. All protocols with mice were approved by the Pennsylvania State University Institutional Animal Care and Use Committee (IACUC protocol # 42678), and experiments conformed to the Association for Assessment and Accreditation of Laboratory Animal Care (AAALAC) guidelines. Adherence to the principles of the ARRIVE guidelines was maintained throughout the study.

### Generation and Validation of Transgenic Parasite Lines

Transgenic *Plasmodium yoelii* (17XNL strain) parasite lines were created using conventional reverse genetics approaches with specific targeting regions for double homologous recombination for each tagged line (79). Targeting regions for the coding sequence (CDS) and 3’ UTR of target genes were PCR amplified from wild-type Py17XNL genomic DNA and combined into a single amplicon by sequence overlap extension (SOE) PCR. The oligonucleotides used in this study to generate and validate transgenic parasite lines are listed in Table S7. This amplicon was inserted into an intermediate plasmid (pCR-Blunt) for initial sequence verification of the target regions. The targeting regions were then inserted into a final pDEF plasmid vector containing the coding sequence for GFPmut2, or TurboID and GFPmut2, with the *P. berghei* DHFR 3’ UTR, and the human DHFR drug resistance cassette (20, 40, 80).

The final plasmids were linearized and transfected into Accudenz-purified *P. yoelii* schizonts produced in *ex vivo* cultures, as previously described (20, 40, 79-82). Schizonts were transfected with 5-10 ug linearized plasmid using an Amaxa Nucleofector 2b device using cytomix with Program T-016. Electroporated cells were injected into the rodent host by tail vein injection, and parasites were selected by pyrimethamine drug cycling in the initially injected mouse (“parental”). Pyrimethamine (ICN Biomedicals Inc., Cat# 194180) was dissolved to 281.4 µM in the mouse drinking water and was provided to the parental mice *ad libitum* starting one day after transfection. The parental mice were provided pyrimethamine drinking water for three days, then were cycled off to regular (no drug) drinking water for at least three days until the parasites were transferred to a second mouse (“transfer”) after reaching 1% parasitemia in the parental mouse. Parasites in the transfer mouse were similarly pyrimethamine cycled to select for transgenic parasites, and parasites were collected by cardiac puncture when infection reached 1% parasitemia. Long-term freezer stocks of collected parasites were prepared by mixing 100 uL of collected blood with 400 uL Freezing Solution (9:1 Alsever’s solution (Sigma-Aldrich, Cat#A3551):glycerol), and stored in liquid nitrogen. Parasite genomic DNA was purified from the blood of a transfer mouse using the Qiagen QIAamp DNA Blood Kit (Qiagen, Cat# 51106), and genotyping PCR was performed to assess integration into the target locus. In some cases, transgenic parasite populations were enriched over remaining wild-type parasites using FACS to select fluorescent parasites.

### Enrichment of Female Gametocytes and Zygotes by Flow Cytometry

Transgenic parasites that express GFP in a stage-enriched manner in female gametocytes or zygotes, respectively, were used for flow cytometry-based enrichment of these parasite stages. A parasite line driving GFP with a female-enriched promoter, *pylap4*, from a dispensable genomic locus (*p230p*) was used to generate the female-enriched samples (22). Blood from mice infected with this *pylap4* prom-GFP parasite line was collected by cardiac puncture as the infection reached 1-3% parasitemia, as measured by Giemsa-stained thin blood smear. The collected blood was maintained at 37°C to prevent gametogenesis, and infected red blood cells were enriched by Accudenz gradient, as described below. Cells collected from the interface of the gradient were filtered before FACS collection of GFP-positive cells on the Beckman Coulter MoFlo Astrios EQ Cell Sorter. Uninfected blood was used as a background fluorescence control. The collected parasites were also maintained at 37°C before they were combined and pelleted at 200 *xg* for 10 minutes at 37 °C or 1034 *xg* for 20 minutes at 37°C. From each replicate sort, between 4 and 8 million female gametocytes were collected and stored in 2 million cell aliquots at -80°C for downstream RNA isolation and total proteomic analyses.

Zygotes were enriched using a transgenic parasite line expressing PyApiAP2-O::GFP and staining with a zygote-specific antibody, anti-Pys25 (Mm, Mab). Mice infected with PyApiAP2-O::GFP tagged parasites were euthanized as they reached 1-3% parasitemia, and blood was collected and maintained at 37°C to prevent premature gametogenesis. Zygotes were cultured as described below, and in the final two hours of zygote culturing, were incubated with anti-Pys25 primary antibody for one hour, then Goat anti-Mouse IgG Alexa Fluor 594 (Invitrogen, A11005) for the final hour. Nuclei were stained with DRAQ5 (Thermo Scientific, Cat # 62251) before cells were pelleted and filtered for sorting on the Beckman Coulter MoFlo Astrios EQ Cell Sorter for Pys25-positive and DRAQ5+ cells. Uninfected blood was used as a background fluorescence control. For each replicate sort, between 4 and 8 million zygotes were collected and stored in 2 million cell aliquots at -80°C for downstream RNA isolation and total proteomic analyses.

### RNA-seq Sample Preparation and Data Analysis

RNA from FACS-collected female gametocytes or zygotes was prepared using the Qiagen RNeasy Kit (Qiagen, Cat#79254) with in-solution DNaseI treatment (Sigma-Aldrich, Cat# AMPD1-KT). RNA quality was assessed by NanoDrop, RNA quantity was determined by Qubit, and RNA integrity was measured by Bioanalyzer. Samples were used to create barcoded libraries using the Illumina Stranded mRNA kit, and an equimolar pool of the samples was sequenced on an Illumina NextSeq to yield 150 nt long single-end reads for each biological replicate. Data was trimmed to remove the low-quality bases and adapter sequence using fastp (0.22.0). Reads shorter than 35 bp after trimming were filtered out. Data quality was inspected by generating FastQC reports before and after trimming. The trimmed reads were mapped to the *P. yoelii* 17X strain reference genome (Plasmodb.org, release 55) using hisat2, version 2.1.0 (83) specifying ’--rna-strandness R’ and ’--max-intronlen 1500’. Coverage files were generated, and mapping was visualized in Integrative Genomics Viewer (IGV). Reads mapping to the genes were counted with featureCounts (version 2.0.1) (84). Parameters ’-t exon -s 2 -g gene_id’ were specified for featureCounts. The Transcript Integrity Number (TIN) was calculated to check the integrity of the transcripts as previously described (40). A complete description of commands used in this analysis is provided in the Makefile (File S1). RNA-seq files are available through NCBI GEO (GSE231838; GSM7304682-GSM7304686).

Read counts were normalized and differential expression between female gametocytes and zygotes was determined using DESeq2 package in R (85). Reads were filtered for quality by transcript integrity number (TIN) to ensure reads have even coverage across the entire length of the gene (40, 86). For filtering, significantly expressed genes with FDR <0.05 and an average TIN <40 in both conditions but with an average count >20 in either of the conditions were flagged for further inspection. TIN-log2-fold change was calculated, and the flagged genes were resolved and included in the differential expression set if the absolute value of TIN log2-FC is < 1.5. The average read counts for transcripts that meet the TIN cut-off were used to calculate the fold change for each transcript detected (average count zygote / average count gametocyte). Transcripts with log2(fold change) > 2.3 were considered to be differentially expressed in zygotes, and those with log2(fold change) < -2.3 were considered differentially expressed in gametocytes. Transcripts detected in either stage were ranked in abundance based on average read count. Heatmaps were generated using DEseq2 normalized counts for differentially expressed genes that were converted to a z-score to denote how far the values deviate from the mean. A positive value (shown in red) indicates upregulation in zygotes, and a negative value (shown in green) indicates downregulation in zygotes. A PCA plot was generated with the plotPCA function available in DESeq2 package, using VST with transformed raw counts as the input.

### Proteomics Sample Preparation

From each of the six gametocyte and zygote biological replicates analyzed by transcriptomics, an aliquot containing 2 million cells was taken for proteomics. One additional sample of 6M cells of zygotes or gametocytes was prepared for the purpose of creating sample-specific spectral libraries. The cells were washed three times with incomplete RPMI, and the pellet in ∼30 µL of media was stored at -80°C.

Sample processing for proteomics was carried out using S-Traps (ProtiFi) (87). Unless otherwise noted, all solid reagents were from Merck-Sigma; solvents and acids were Optima LC-MS grade from Fisher Scientific. Water was LC-MS grade from Honeywell Burdick & Jackson. To each cell pellet of ∼30 µL was added 30 µL of 10% w/v sodium dodecyl sulfate in 100 mM ammonium bicarbonate (ABC) and 2.6 µL 120 mM tris(2-carboxyethyl)phosphine (TCEP, Pierce Bond Breaker) in 50 mM ABC. Samples were incubated for 5 min in a 95°C water bath with occasional vortexing, after which they were cooled to room temperature and centrifuged 2 min at 20,000 *× g*. The volume was estimated by pipetting and additional water was added to achieve a final volume of 62.6 µL, then 2.6 µL of 0.5 M iodoacetamide in 50 mM ABC was added, and the samples were incubated for 20 min in darkness at room temperature with vortexing. Samples were acidified by adding 6.5 µL of 27.5 % v/v phosphoric acid and the protein was precipitated by adding 430 µL of S-Trap buffer (90% v/v methanol, 100 mM Tris pH 7.0). Protein was loaded onto S-Traps by adding precipitate suspension in 100 µL increments and centrifuging 30-60s at 1500 *× g.* The traps were washed three times with 150 µL S-Trap buffer and placed in clean 1.5 mL microcentrifuge tubes (Eppendorf Protein LoBind). Then 20 µL of 50 mM ABC containing 1 µg of trypsin (Promega Platinum Mass Spectrometry Grade) was added and the traps were incubated 2h at 47°C in a covered thermomixer without vortexing. After digestion, 40 µL of water was added to the traps and peptides were recovered by centrifuging. Two additional recovery washes were performed and combined with the first wash: 40 µL of 0.2% formic acid (FA) and 30 µL 50% acetonitrile (ACN). The combined eluates were dried in a vacuum concentrator and resuspended in 20 µL 0.1 % trifluoroacetic acid (TFA).

### LC-MS

LC was performed with an EASY-nLC 1000 (Thermo Fisher Scientific, USA) using a vented trap set-up. The trap column was a PepMap 100 C18 (Thermo Fisher Scientific #164946) with 75 µm ID and a 2 cm bed of 3µm 100 Å C18. The analytical column was an EASY-Spray column (ThermoFisher Scientific #ES804) with 75 µm ID and a 15 cm bed of 2µm 100 Å C18 operated at 35°C. The LC mobile phases (Honeywell Burdick & Jackson) consisted of buffer A (0.1 % v/v formic acid in water) and buffer B (0.1 % v/v formic acid in ACN). The separation gradient, operated at 300 nL/min, was 4 % B to 28 % B over 85 min, 28% B to 30% B over 5 min, 30% B to 80% B over 5 min, and 10 min at 80% B. Prior to each run, the trap was pre-conditioned with 5 µL buffer A at 800 bar, the column was preconditioned with 3.5 µL buffer A at 800 bar, and the sample was loaded onto the trap with 15 µL buffer A. DIA-MS was performed using a Thermo Fisher Scientific Orbitrap Eclipse. The six biological replicate samples were analyzed by a tSIM method with the following acquisition settings: MS1 scan from 395-1005 *m/z* at 30,000 resolution, default AGC settings (target of 4×10^5^ ions, max ion injection time set to “Auto”); MS2 at 15,000 resolution with an AGC target of 1000% (5×10^5^ ions) and a maximum injection time of 22 ms, HCD fragmentation at 33% normalized collision energy, 8 *m/z* isolation window, and a loop count of 75 with an inclusion list of 151 overlapping DIA windows with optimized *m/z* values generated using the tool in EncyclopeDIA. The library samples were analyzed using gas-phase fractionation (28), a method wherein the same sample was injected six times with each method narrowed to a 100 *m/z*-wide band of masses, i.e., 395-505, 495-605…895-1005. The DIA-MS methods were the same as above except for the following: MS1 scan range was narrowed to 100 *m/z*; MS2 maximum injection time of 54 ms, 4 *m/z* isolation window, and a loop count of 25 with an inclusion list of 51 overlapping DIA windows.

### DIA data analysis

Raw mass spectrometry data were converted to mzML using msConvert version 3.0.19106 (Proteowizard (88)). Peaks were centroided and the staggered DIA windows were demultiplexed (29). DIA data were analyzed with EncyclopeDIA version 1.12.31 (29) using Open JDK 13 on a Slurm 19.05.5 cluster running under Ubuntu 20.04. Default parameters were used except the following: Percolator version 3-01 was used, Percolator training set size was increased to 1M, and the -quantifyAcrossSamples flag was set to true in all cases except when exporting chromatogram libraries. Spectral libraries were generated *in silico* using Prosit (27) as described previously (26). Briefly, the reference *P. yoelii* 17X (89) protein FASTA database (PlasmoDB, version 55) and a mouse database (Uniref90 (90) obtained from UniProt.org (91) were digested *in silico* using the tool in EncyclopeDIA (26) to obtain a list of all theoretical peptides with *m/z* between 396.4 and 1004.7, two tryptic termini, up to one missed cleavage, and charge state 2 or 3. This list was uploaded to Prosit (www.proteomicsdb.org/prosit/) and theoretical spectra were generated for the peptides using the 2020 HCD intensity prediction model the 2019 iRT retention time prediction model, assuming an NCE of 33 and a default charge state of 3. The library samples (comprising six gas-phase fractions) were searched against this *in silico* library, and identified peptides were mapped to a protein database comprising the *P. yoelii* and mouse protein databases and contaminants (www.thegpm.org/crap/) in order to identify degenerate peptides that could theoretically come from host and contaminant proteins. The identified peptides were exported as sample-specific chromatogram libraries, one for gametocytes and one for zygotes. The DIA-MS data from the biological replicates (three each for gametocytes and zygotes) were searched against their respective chromatogram libraries, and separate protein quantifications were carried out for the two sample types.

### Proteomic Data Processing

Normalization and processing of the raw protein areas was performed in Microsoft Excel (Tables S2, S3). The protein areas of the six biological replicates were log-transformed to produce normal distributions. Correction factors for each sample were derived from the subset of protein areas quantified in all six samples by using z-score normalization to give the subset populations the same mean and standard deviation:

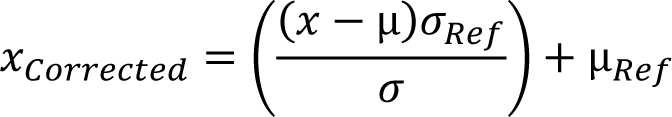

where 𝑥 is the area of a given protein in a sample, µ and 𝜎 are the average and standard deviation, respectively, of the protein areas in the sample used for normalization, and *Ref* is the reference sample. After normalization, the average protein area for a given protein in gametocytes or zygotes was calculated as the geometric mean, i.e., the average of the log-transformed, normalized areas. Where the protein was quantified in at least two of three replicates for each sample type, a p-value was calculated using a two-tailed, homoscedastic t-test. The Benjamini-Hochberg test estimated a false discovery rate (FDR) of less than 8% at a p-value of 0.05. Protein abundances were considered significantly different between gametocytes and zygotes if they were greater than five-fold with p<0.05. Additionally, in order to avoid false negatives and spuriously large ratios that can arise near the limit of detection, proteins were required to be above the 25th percentile of abundance to be considered differently regulated.

To extract relative protein abundances from the gas-phase fractionated library samples, a tool in EncyclopeDIA was used to combine the six gas-phase fractions for each sample type into a single data file that could be searched against the Prosit *in silico* library, and a protein quantification report was generated comparing the gametocyte versus zygote library samples. The reported protein area for a given protein in a given sample (gametocytes or zygotes) was only retained if peptides corresponding to that protein were confidently identified (FDR < 1%) in that sample. The protein areas were transformed and normalized by z-score and relative abundance percentiles were calculated as above. Absent sufficient samples to perform a t-test, a p-value for each gametocyte:zygote ratio was assigned using the complementary error function (erfc). The population of log-transformed gametocyte:zygote protein ratios (i.e., the difference of the log-transformed and normalized protein areas) produced a normal distribution with a mean near zero, i.e., a gametocyte:zygote protein abundance ratio of 1:1. A Gaussian curve with the equation:

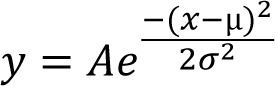

where 𝐴 is the curve height, µ is the mean, and 𝜎 is the standard deviation, was fitted to the population using the solver function in Excel. The values of µ and 𝜎 were used to calculate a p-value for each ratio 𝑥 using the complementary error function ERFC:

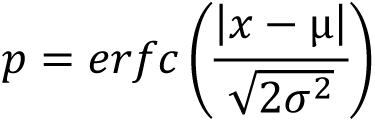

An erfc-derived p-value <0.05 corresponds to a ratio whose magnitude is greater than 95% of the ratios in the population, i.e., the tails of the distribution.

### Searching DIA Data for Post-Translational Modifications

Deconvoluted DIA mzML files were converted into pseudo-DDA spectra using the DISCO tool in the Trans-Proteomic Pipeline (TPP) version 6.2.0 (75). The resulting spectra were searched against the combined *P. yoelii*, mouse, and contaminant FASTA database using Comet version 2020.01 rev. 3 (92). Semi-tryptic peptides with up to two missed cleavages were allowed, with variable modifications of 15.9949 at Met for oxidation, 79.966331 at Ser and Thr for phosphorylation, and 42.010565 at the protein N-terminus or the next residue (in case of cleavage of N-terminal Met) for acetylation. The four experiments (biological replicate and library samples of gametocytes and zygotes) were analyzed separately in the TPP using PeptideProphet and iProphet. Spectra with PeptideProphet scores >0.9 and iProphet scores >0.8 were retained. Although the decoy-estimated, spectrum-level FDR rate at these thresholds was <0.1%, the FDR among putatively phosphorylated spectra was ∼3% owing to the fact that spurious assignment of PTMs is likely to be much higher for an un-enriched sample. Only spectra with iProphet scores corresponding to a decoy-estimated FDR <1% among putative phosphopeptide spectra were retained. (Table S6).

### Enrichment and Preparation of Gametocytes or Zygotes for TurboID Proteomics

To generate gametocyte samples for proximity proteomics, mice were injected with transgenic ALBA4::TurboID::GFP, DOZI::TurboID::GFP or the TurboID::GFP control parasite line, and as the infections reached 1% parasitemia, the mice were provided with water supplemented with 10 mg/L sulfadiazine (VWR, Cat# AAA12370-30) for two days. Gametocytes were selected with sulfadiazine for two days before blood was collected by cardiac puncture for *ex vivo* supplementation with 150 µM biotin in RPMI 1640, 20% v/v FBS at 37°C for 1 hour. After 1 hour, parasites were washed in 1x PBS three times before lysis in a modified RIPA lysis buffer (50 mM Tris-HCl (pH 8.0 at room temperature), 0.1% w/v SDS, 1 mM EDTA, 150 mM NaCl, 1% v/v NP40, 0.5% w/v sodium deoxycholate) with a 1x protease inhibitor cocktail (Roche, VWR, Cat# PI88266) and 0.5% v/v SUPERase In (Life Technologies, Cat# AM2694) at 4°C with end-over-end rotation.

To generate zygote samples for proximity proteomics, mice were infected with the same TurboID transgenic parasite lines as above but were not treated with sulfadiazine. When parasitemia reached 1%, blood was collected by cardiac puncture into incomplete RPMI with 25mM HEPES and L-Glutamine (VWR Cat# 45000-412) at 37°C and enriched by an Accudenz discontinuous gradient as described above. Briefly, the Accudenz collected cells were added to *in vitro* ookinete media (RPMI 1640, 20% v/v FBS (Fisher Scientific, Cat#350-11-CV Lot # 35011126), 0.05% w/v hypoxanthine (Fisher Scientific, Cat#AC122010250), 100 μM xanthurenic acid (Sigma Aldrich, Cat# D120804-1G), pH 8.2 at 22°C) and were allowed to develop for 6 hours at room temperature, to allow for fertilization to occur and to enable zygote development (93-95). Zygotes were observed by DIC and fluorescence microscopy (Zeiss Axioscope A1 with 8-bit AxioCam ICc1 camera) using the 100X oil objective and processed by Zen 2012 (blue edition) imaging software. In the final hour of zygote culture, the parasites were supplemented with 150 µM biotin to allow for biotinylation to occur. After one hour, cells were washed in 1X PBS, and added to magnetic Protein G beads (Invitrogen, Cat# 10003D) coated with mouse monoclonal anti-Pys25 antibody (96) to enrich for zygotes that express Pys25 on their surface away from female gametocytes that do not. The enriched zygotes were lysed on-bead with the above modified RIPA lysis buffer at 4°C for one hour with end-over-end rotation.

The gametocyte and zygote lysates were mechanically lysed with a 1 mL Dounce homogenizer (Wheaton, Cat# 357538) for one minute on ice with a tight pestle following chemical lysis. The parasite lysates were added to magnetic Dynabeads MyOne Streptavidin T1 (Life Technologies, Cat# 65601) for three hours at 4°C with end-over-end rotation to allow the streptavidin beads to bind the biotinylated proteins. The beads were washed with ‘wash buffer’ (50 mM Tris-HCl (pH 8.0 at room temperature), 1 mM EDTA, 150 mM NaCl, 1% v/v NP40) once, and then transferred to a new tube to reduce background from non-specific proteins binding to the surface of the plastic tube. The beads were washed three more times with this wash buffer, then resuspended in 1x PBS. A quarter of the beads were resuspended in 2X Sample Buffer (50 mM Tris-HCl (pH 6.8 at RT), 5% w/v SDS, 5% v/v glycerol, 0.16% w/v bromophenol blue, 200 mM NaCl) with a fresh addition of 5% v/v β-mercaptoethanol and 1 mM biotin. Samples were then heated to 95°C to denature streptavidin and elute the biotinylated proteins off the beads. The eluted proteins were removed from the beads and used for affinity blot analyses, probed with anti-GFPmut2 primary antibody (1:1000 dilution, rabbit pAb, custom Pocono RF&L) with anti-rabbit conjugated to HRP (1:1000 dilution, Invitrogen, Cat #A16104) or streptavidin-HRP (1:1000 dilution, Fisher Scientific, Novex Cat# 43-423-3) to ensure that biotinylated and bait proteins were present. Affinity blots were imaged with Pierce™ ECL Western Blotting Substrate solution (Thermo Scientific, Cat# 32106) and imaged with BioRad Gel Dox XR Imaging System.

### Tandem Mass Tag (TMT) Mass Spectrometry for the Identification of TurboID Proximal Proteins

The on-bead captured TurboID samples were analyzed by the Indiana University School of Medicine Proteomics Core. The samples were digested on-bead with LysC/Trypsin in 8M urea, followed by a peptide clean-up on SPE spin columns before TMT labeling with the TMTpro™ 16plex Label Reagent Set (Thermo Fisher, Cat# A44520). Peptides from each sample were labeled with a distinct TMTpro™ isobaric tag before mixing. Samples were combined by sample type (e.g., unfused control replicates, ALBA4 experimental replicates, DOZI experimental replicates) in equal volumes. Each mix was then run on the Exploris Orbitrap mass spectrometer with 3 FAIMS (field asymmetric ion mobility spectrometry). Peptides were mapped to a reference database of *P. yoelii* PY17X annotated proteins (plasmodb.org release 48) (23), *Mus musculus* proteome (UniProt Proteome ID: UP000000589) (97), and the Contaminant Repository for Affinity Purification (CRAPome 2.0) using Proteome Discoverer, version 2.5 to determine protein abundance values for each identified protein (98). The PY17X proteins were filtered, and the average abundance values for the experimental samples and control were compared. The abundance ratio was calculated to determine the enrichment of proteins detected in experimental over control (abundance experimental / abundance control). A log2(abundance ratio) >2 was considered enriched in the experimental sample over the control.

### IFA Sample Preparation and Structured Illumination Microscopy

PyDOZI::GFP and PyNOT1-G::GFP expressing female gametocytes were imaged by indirect immunofluorescence microscopy on the BioVision Technologies VT-iSIM super-resolution microscope on 100X (oil) objective for high-resolution image capture of Z-slices every 1µm with MetaMorph software. Images were deconvolved using the Microvolution plug-in on ImageJ, and colocalization was measured using the JACoP ImageJ plug-in (72).

Cells were prepared for IFA as previously described (22, 40). Briefly, the transgenic gametocytes were pelleted at 1,400 *xg* for 3 minutes and fixed in 4% v/v paraformaldehyde (VWR, Cat# PI28908) and 0.0075% v/v glutaraldehyde (VWR, Cat#AAAA17876-AE) in 1xPBS for with rocking for 30 minutes at room temperature. Cells were washed once in 1x PBS and permeabilized in 1% v/v Triton X-100 (Fisher Scientific, Cat # AC327372500) for 10 minutes at room temperature. Cells were well washed with 1x PBS to remove permeabilization solution before blocking with 3% w/v BSA (Sigma-Aldrich, Cat # A7906-100G) in 1x PBS overnight at 4°C. Blocked cells were stained with primary antibody at 1:1000 dilutions in the 3% w/v BSA blocking solution for one hour at room temperature with rocking. Primary antibodies used were against GFP (Mouse mAb 4C9, DSHB, Cat# DSHB-GFP-4C9), HsDDX6 (DOZI homolog, rabbit pAb) (20, 80), or PyPABP1 (rabbit pAb, Pocono RF&L, custom antibody) (71). Cells were washed after incubation with the primary antibody twice with 3% w/v BSA blocking solution, then resuspended in the blocking solution and incubated with the secondary antibodies for 1 hour, all at 1:1000 dilutions. Secondary antibodies used were against mouse (Donkey anti-mouse, AF488; Invitrogen Cat#A21202) or goat (Donkey anti-goat AF488, Invitrogen Cat# A11055) or rabbit (Donkey anti-rabbit AF594, Invitrogen, Cat# A11012). Cells were washed in 1x PBS, then stained with 1 µg/mL DAPI (4′,6-diamidino-2-phenylindole) in 1x PBS for 5 min at room temperature. Cells were washed once in 1x PBS, then resuspended in 1xPBS before mounting on glass slides in a 1:1 ratio with ProLong Gold Antifade Mountant (Invitrogen, Cat# P36930), and covered with a glass cover slip before imaging.

### Ultrastructure Expansion Microscopy (U-ExM)

PyDOZI::GFP expressing female gametocytes were subjected to ultrastructure expansion microscopy (U-ExM) as previously described (68, 99). Cells were stained with chicken anti-GFP IgY fraction (Aves Labs, #GFP-1010) at a 1:1000 dilution, and counterstained with custom rabbit polyclonal antisera against PyCITH, PyPABP1, or PyALBA4 (20, 70, 71) at a 1:5000 dilution. Secondary antibodies against chicken IgY (Thermo, #A11039, AF488) or rabbit IgG (Thermo #A11011, AF568) were applied at a 1:500 dilution. Sytox Deep Red (Thermo, #S11380, 1:1000) and NHS-Ester (Thermo, #30000, AF405, 1:250) were used to stain nucleic acids and proteins as before (68, 99). Images were captured on a Zeiss LSM980 series microscope using a Plan-Apochromat 63X/1.40 Oil DIC M27 objective with Airyscan 2 detector using ZEN Blue software. Z-stacks were acquired at zoom 5.0 with a 0.1 um step size. Images were processed using ZEN AiryScan Joint Deconvolution Module followed by channel alignment.

## Data Availability

Datasets associated with this study are publicly available: RNA-seq files are available at NCBI GEO: SRA BioProject: GSE231838; GSM7304682-GSM7304686. The mass spectrometry data associated with this work are available through the MassIVE repository (https://massive.ucsd.edu/) with the identifier MSV000092154 or ProteomeXchange (http://www.proteomexchange.org) with the identifier PXD042955.

## Acknowledgments

The authors would like to acknowledge the Huck Institutes’ Core Facilities of Penn State at University Park that were instrumental in conducting this work (Genomics: RRID:SCR_023645; Microscopy: RRID: SCR_024457). We also thank Amber Mosley and Emma Doud of the Indiana University School of Medicine Proteomics Core for their expert assistance in generating the tandem mass tag proteomic data, and for critical discussions of the resulting data. We are grateful to Michael Tribone of the Huck Institutes for assistance in creating the model illustration (Fig 6). All of our work is greatly enabled and impacted by VEuPathDB as an essential resource. We thank Takafumi Tsuboi for the provision of the anti-Pys25 hybridoma. Finally, we also acknowledge members of the Llinás and Lindner laboratories for critical discussions of this work.

## Author Contributions: (CREDIT Designations)

Conceptualization: KTR, SEL

Data curation: KTR, AS, MF, KES, SEL

Formal analysis: KTR, AS, KES, SEL

Funding acquisition: AS, RLM, KES, SEL

Investigation: KTR, JPM, KES, SEL

Methodology: KTR, SA, MF, KES, SEL

Project administration: KTR, SEL

Resources: MF, RLM, KES, SEL

Software: RLM, KES

Supervision: SEL

Validation: KTR, KES

Visualization: KTR, JPM, KES, SEL

Writing – original draft: KTR, KES, SEL

Writing – reviewing and editing: KTR, JPM, RLM, MF, SA, KES, SEL

## Supp Figures

**Figure S1: Genomic locus and genotyping PCR for the PyApiAP2-O::GFP lines.** (Top) A genomic locus schematic of the unedited wild-type locus and the transgenic locus are shown with anticipated PCR fragment sizes listed. (Bottom) Genotyping PCR results for wild-type (Py17XNL), transgenic parasites (TG), no template controls (NTC), and a positive plasmid control (the plasmid used to create the transgenic line) are shown.

**Figure S2: Flow cytometric enrichment of female gametocytes and zygotes.** A. Uninfected red blood cells were used as a negative control for the collection of female gametocytes expressing green fluorescent protein (GFP) from a female-enriched promoter. B. Parasites expressing GFP from the female-enriched *pylap4*_promoter_ were selected by fluorescence activated cell sorting and collected into RPMI kept at 37°C to prevent gametogenesis. C. Collected cells were assessed by fluorescence microscopy to ensure female gametocytes were specifically enriched by FACS. D. Uninfected red blood cells stained with DRAQ5 nuclear stain were used as a negative control for the collection of Pys25-positive zygotes. E. *In vitro* cultured zygotes were surface stained with α-Pys25 primary antibody (mouse) and α-mouse Alexa Fluor594 secondary antibody, and DRAQ5 to separate the Pys25-positive zygotes from uninfected red blood cells or other parasite life stages. The cultured zygotes also expressed GFP as a fusion protein with PyApiAP2-O::GFP. F. Collected zygotes were assessed by live fluorescence to ensure the expected population was enriched by FACS.

**Figure S3: Genomic locus and genotyping PCRs for *P. yoelii* Parasites with TurboID-GFP Fusions.** Genomic locus schematics and validation data for (A) the unfused control, (B) PyALBA4, and (C) PyDOZI are provided. (Top) The unedited wild-type locus and the transgenic locus are illustrated with anticipated PCR fragment sizes listed. (Bottom) Genotyping PCR results for wild-type (Py17XNL), transgenic parasites (TG), no template controls (NTC), and a positive plasmid control (the plasmid used to create the transgenic line) are shown.

**Figure S4: Quality control blots in support of TurboID experiments with female gametocytes and zygotes.** A. Schematic of TurboID::GFP-tagged PyALBA4, PyDOZI, and an unfused control. Expected protein masses of each fusion protein are provided. (B and C) PyALBA4 was endogenously tagged with GFP (no TurboID), BioID::GFP, or TurboID::GFP (TID). Mixed blood stage parasites were supplemented without (0) or with 150uM biotin for 15 minutes (15’) or 12 hours. Whole-cell lysates were probed with (B) streptavidin-HRP to assess the extent of biotinylation in each sample or (C) α-GFP antibody to confirm that while the efficiency of biotinylation differs between these parasite lines, the tagged proteins are present at qualitatively similar abundances. (D and E) Gametocytes were similarly tested for TurboID activity in an *ex vivo* culture without biotin supplementation (0) or with 150uM biotin for 15 minutes, or 1, 2, 4, or 12 hours. Unfused TID::GFP gametocytes were compared with (D) PyALBA4::TID::GFP or (E) PyDOZI::TID:GFP. Whole-cell lysates were probed with streptavidin-HRP to assess the extent of biotinylation in each sample. A red asterisk at the bottom of the 1-hour lane indicates that this condition was selected for mass spectrometric analyses. (F) Zygotes were similarly tested for TurboID activity. Zygotes were cultured *in vitro* for 6 hours, with or without supplementation with 150 μM biotin for the final hour before capture on Pys25-coated magnetic Protein G beads. Unfused TID::GFP zygotes were compared with PyALBA4::TID::GFP or PyDOZI::TID::GFP. Whole-cell lysates were probed with streptavidin-HRP to assess the extent of biotinylation in each sample. (G and H) TurboID-based biotinylated proteins from gametocytes expressing unfused TID::GFP or PyALBA4::TID::GFP were captured on streptavidin-conjugated Dynabeads. The input (“I”), flow-through (“F”), and eluate (“E”) were probed with (F) streptavidin-HRP or (G) α-GFP antibody as above. (I and J) The same experiment but with TID::GFP and PyDOZI::TID::GFP was conducted and probed as in panels F and G. (K) TurboID-based biotinylated proteins from *in vitro* zygotes expressing unfused TID::GFP, PyALBA4::TID::GFP, or PyDOZI::TID::GFP were captured on streptavidin-conjugated Dynabeads. The input (“I”), flow-through (“F”), and eluate (“E”) were probed with (F) streptavidin-HRP.

**Figure S5: Comparisons to previously identified protein interactions of DOZI and ALBA4.** (A) Proteins proximal to PyDOZI::TurboID::GFP and/or PyALBA4::TurboID::GFP in gametocytes were compared to the immunoprecipitated PbDOZI::GFP complex in gametocytes by Mair and colleagues (10). (B) Proteins proximal to PyALBA4::TurboID::GFP in gametocytes were compared to the immunoprecipitated PyALBA4::GFP complex in gametocytes by Munoz and colleagues (20). (C) Proteins proximal to PyDOZI::TurboID::GFP in gametocytes were compared to the immunoprecipitated PyCCR4-1::GFP complex in blood stages by Hart and colleagues (80).

**Figure S6: Colocalization analysis of detected protein-protein interactions in female gametocytes**. PyDOZI::GFP-expressing (A) or PyNOT-1G::GFP-expressing (B) female gametocytes were used for super-resolution structured illumination microscopy (3D-SIM) imaging to assess protein colocalization. The ImageJ plugin JACoP was used to assess 3D fluorescence colocalization for each pair of proteins indicated (72, 100). The signal intensity from the green channel was compared to the corresponding pixel in the red channel for all Z-stacks of the 3D image to calculate the overlap coefficient. This value varies from 0 to 1, with 0 indicating no fluorescence signal overlap and 1 reflecting complete fluorescence signal colocalization.

## Supp Tables

Table S1: Complete datasets for RNA-seq experiments completed in this study.

Table S2: Complete datasets for DIA proteomics of spectral library samples completed in this study. Table S3: Complete datasets for DIA proteomics of experimental samples completed in this study.

Table S4: Translationally repressed mRNAs across the host-to-vector transmission event with comparisons to previous datasets (Guerreiro *et al*. (21), Lasonder *et al*. (6), Sebastian *et al*. (74)).

Table S5: Complete datasets of TurboID-based proximity proteomics of PyDOZI and PyALBA4 in female gametocytes and zygotes.

Table S6: Complete dataset of post-translational modifications detected by mass spectrometry.

Table S7: Oligonucleotides used in this study. Lower case letters indicate non-homologous bases added for cloning purposes.

## Supp Files

File S1: A Makefile describing the bioinformatic workflow used for RNA-seq analyses in this study.

