## Supplementary material for "Global Release of Translational Repression Across *Plasmodium’s* Host-to-Vector Transmission Event": File S1: deseq2.r

```
#!/usr/bin/env Rscript
#
# The program creates differential expression results using the deseq2 method.
#
# Usage:
#
# cat counts.txt | Rscript deseq2.r NxM
#
# Produces a CSV file table with differentially expressed rows printed to the standard output.
#
# The input count file is assumed to contain the count data in the last N + M columns.
#
# The NxM represents the experimental design of columns: 3x3
#
# Assumes the first column is the feature name and 
# the last N + M columns of are the count matrix.
#

# Command line argument.
args = commandArgs(trailingOnly=TRUE)  

# Input stream.
infile = "stdin"

# Output stream.
outfile = ""

if (length(args)!=1) {
  stop("Experimental design must be specified as: NxM at the command line", call.=FALSE)
}

# Load the library while suppressing verbose messages.
suppressPackageStartupMessages(library(DESeq2))

# The first argument to the script.
first = args[1]

# Extract the experimental design from the command line.
design = unlist(strsplit(first, 'x'))

# Find the desing counts.
reps1 = as.integer(design[1])
reps2 = as.integer(design[2])

# Set up the conditions based on the experimental setup.
cond1 = rep("cond1", reps1)
cond2 = rep("cond2", reps2)

# Read the data from the standard input.
counts = read.table(infile, header=TRUE, sep="\t", row.names=1 )

# Assume the last N + M columns are the count matrix.
idx = ncol(counts) - (reps1 + reps2)

# Cut out the valid columns.
#counts = counts[-c(1:idx)]
if (idx > 0) counts = counts[-c(1:idx)] else counts=counts

# Some tools generate the estimated counts as real numbers
# DESeq 2 allows only integers. We need to convert real numbers to rounded integers.
numeric_idx = sapply(counts, mode) == 'numeric'
counts[numeric_idx] = round(counts[numeric_idx], 0)

# Build the dataset
samples = names(counts)
condition = factor(c(cond1, cond2))
colData = data.frame(samples=samples, condition=condition)

# Create DESEq2 dataset.
dds = DESeqDataSetFromMatrix(countData=counts, colData=colData, design = ~condition)

#Set the reference to be compared
dds$condition = relevel(dds$condition,"cond1")

# Run deseq2.
dds = DESeq(dds)

# Extract the DESeq2 results.
res = results(dds)

# Turn the DESeq2 results into data frames.
data = data.frame(res)

# Rename columns for what they are.
names(data)[names(data)=="pvalue"] <-"PValue"
names(data)[names(data)=="padj"] <-"FDR"

# Create the additional columns.
data$baseMeanA = 1
data$baseMeanB = 1
data$foldChange = 2 ^ data$log2FoldChange
data$PAdj = p.adjust(data$PValue, method="hochberg")

# Sort the data by PValue to compute false discovery counts.
data = data[with(data, order(PValue, -foldChange)), ]

# Compute the false discovery counts on the sorted table.
data$falsePos = 1:nrow(data) * data$FDR

# Rearrange the columns into the same order as other methods do.
data = data[c(1,7,8,9,2,3,4,5, 10, 6, 11)]

# Get the normalized counts.
normed = counts(dds, normalized=TRUE)

# Round normalized counts to a single digit.
normed = round(normed, 1)

# Merge the two datasets by row names.
total <- merge(data, normed, by=0)

# Sort again for output.
total = total[with(total, order(PValue, -foldChange)), ]

# Start index for normalized data for condition 1.
start1 = which(colnames(total) == "falsePos") + 1

# End index for normalized data for condition 1.
end1 = start1 + reps1 - 1

# Start index for normalized data for condition 2.
start2 = end1 + 1

# End index for normalized data for condition 2.
end2 = start2 + reps2 - 1

# Compute the totals.
total$baseMeanA = rowMeans(total[,start1: end1])
total$baseMeanB = rowMeans(total[,start2: end2])

# Bringing some sanity to numbers. Rounding some columns to fewer digits.
total$foldChange = signif(total$foldChange, 3)
total$log2FoldChange = round(total$log2FoldChange, 1)
total$baseMean = round(total$baseMean, 1)
total$baseMeanA = round(total$baseMeanA, 1)
total$baseMeanB =  round(total$baseMeanB, 1)
total$lfcSE = round(total$lfcSE, 2)
total$stat = round(total$stat, 2)
total$FDR = signif(total$FDR, 4)
total$falsePos = round(total$falsePos, 0)

# Rename the row name column.
colnames(total)[1] <- "name"

# Write the results to the standard output.
write.csv(total, file=outfile, row.names=FALSE, quote=FALSE)
```
