## Supplementary material for "Global Release of Translational Repression Across *Plasmodium’s* Host-to-Vector Transmission Event": File S1: Makefile

```
SAMPLES=sampleinfo.txt
DATA=data
TRIM=$(DATA)/trim
FASTQC_RAW=$(DATA)/fastqc/raw
FASTQC_TRIM=$(DATA)/fastqc/trim
FASTQC=$(FASTQC_TRIM)

REFS=refs
#GENOME=$(REFS)/PlasmoDB-55_Pyoeliiyoelii17X_Genome.fasta
GENOME=$(REFS)/17XNL_PSU.fa
GFF=$(REFS)/PlasmoDB-55_Pyoeliiyoelii17X.gff
GTF=$(REFS)/PlasmoDB-55_Pyoeliiyoelii17X.gtf


HISAT=hisat2_17XNL_PSU
GCOUNTS=$(HISAT)/gene_counts


fastqc:
	#
	# Generate quality reports
	#
	mkdir -p $(FASTQC)
	cat $(SAMPLES) | parallel --verbose --progress -j 6 fastqc --nogroup -o $(FASTQC) $(TRIM)/{}.fq.gz

multiqc:
	#
	# Generate combined quality report
	#
	multiqc $(FASTQC) -o $(FASTQC)/multiqc_trim

fastp_trim:
	#
	# Trim adapters using fastp
	#
	mkdir -p $(TRIM) $(TRIM)/reports
	cat $(SAMPLES) | parallel --verbose --progress -j 6 fastp -i $(DATA)/{}_R1_001.fastq.gz -o $(TRIM)/{}.fq.gz --cut_right --cut_right_window_size 4 --cut_right_mean_quality 24 -l 35 -f 1 -t 1 -h $(TRIM)/reports/{}.html -j $(TRIM)/reports/{}.json -a CTGTCTCTTATA 

data_stats:
	seqkit stats $(DATA)/*.gz >data_stats_raw.txt
	seqkit stats $(TRIM)/*.gz >data_stats_trim.txt

hisat2_aln:
	mkdir -p $(HISAT)
	cat $(SAMPLES) | parallel --verbose --progress -j 12 'hisat2 -q -x $(GENOME) --rg-id {} --rg SM:{} --rna-strandness R -U $(TRIM)/{}.fq.gz -S $(HISAT)/{}.sam --max-intronlen 1500 2>$(HISAT)/{}_summary.txt'


sam2bam:
	#
	#convert sam to bam
	#
	cat $(SAMPLES) |parallel -j 8 --verbose --progress 'samtools view -b $(HISAT)/{}.sam | samtools sort - -o $(HISAT)/{.}.bam -O bam -T $(HISAT)/{}_tmp'
	ls $(HISAT)/*bam | parallel -j 8 --verbose --progress samtools index {}

coverage:
	ls $(HISAT)/*.bam | parallel -j 6 --progress --verbose 'bedtools genomecov -ibam  {}  -split -bg  > {.}.bedgraph'
	ls $(HISAT)/*.bedgraph |parallel -j 8 --progress --verbose bedSort {} {.}.s.bedgraph
	ls $(HISAT)/*.s.bedgraph |sed 's/.s.bedgraph//' | parallel -j 3 --progress --verbose 'bedGraphToBigWig {}.s.bedgraph  $(GENOME).fai {}.bw'

gtf:
	gffread -T -o $(GTF)  $(GFF)

featurecounts:
	#
	# Count reads at gene level.
	#
	mkdir -p $(GCOUNTS)
	featureCounts -T 8 -t exon -s 2 -g gene_id -a $(GTF) -o $(GCOUNTS)/gcounts.txt $(HISAT)/Fe*.bam $(HISAT)/Zy*.bam
	#
	# format file.
	cat $(GCOUNTS)/gcounts.txt | grep -v "#" | sed -e '1s/.bam//g' -e '1s/hisat2\///g' >$(GCOUNTS)/gene_counts.txt 

tin:
	#
	# Calculate TIN from bam files.
	#
	mkdir -p tin
	tin --ann $(GTF) --feat exon --groupby gene_id --strand reverse --lib_type single --read_len 146 $(HISAT)/Fe*.bam $(HISAT)/Zy*.bam  >tin/gene_tin.txt


overlap:
	#
	# Check if there is overlap in transcription from neighbouring genes.
	#
	mkdir -p tin
	overlap --ann $(GTF) --strand reverse --n 0 --read_len 146 --lib_type single --tin_cutoff 40 --count_cutoff 20 $(HISAT)/Fe*.bam $(HISAT)/Zy*.bam >tin/runin.txt
```
