## Supplementary material for "Global Release of Translational Repression Across *Plasmodium’s* Host-to-Vector Transmission Event": File S1: README.txt

Scripts used in the RNAseq analysis of female gametocytes and zygotes

1. Makefile - Makefile with all commands used in the bioinformatics analysis.
2. deseq2.r - Custom R script used for DESeq2 differential expression analysis.
3. heatmap.r - Custom R script used to generate DEG heatmaps.
4. summary_plots.r - Custom R script used in the generation of pca plot.

Note:

Transcript Integrity Number (TIN) was calculated using the code from the github repository  https://github.com/aswathyseb/tincheck
