## Supplementary material for "Global Release of Translational Repression Across *Plasmodium’s* Host-to-Vector Transmission Event": File S1: summary_plots.r

```
#!/usr/bin/env Rscript
#
# This script uses function in DESeq2 package and  generates pca plot and sample-to-sample distance heatmap of all samples.
#
# Usage:
# cat counts_mod.txt | Rscript summary_plots.r
#

# Set the plot dimensions.
WIDTH = 10
HEIGHT = 5

# Input stream.
infile = "stdin"

reps1 = 2
reps2 = 3

# Load the library while suppressing verbose messages.
suppressPackageStartupMessages(library(DESeq2))
suppressPackageStartupMessages(library(ggplot2))

# Set up the conditions based on the experimental setup.
gametocytes = rep("Fe", 2)
zygotes = rep("Zy", 3)

cond = c(gametocytes,zygotes)

# Read the data from the standard input.
counts = read.table(infile, header=TRUE, sep="\t", row.names=1 )

# Assume the last N + M columns are the count matrix.
idx = ncol(counts) - (reps1 + reps2 )

# Cut out the valid columns.
#counts = counts[-c(1:idx)]
if (idx > 0) counts = counts[-c(1:idx)] else counts=counts

# Some tools generate the estimated counts as real numbers
# DESeq 2 allows only integers. We need to convert real numbers to rounded integers.
numeric_idx = sapply(counts, mode) == 'numeric'
counts[numeric_idx] = round(counts[numeric_idx], 0)

# Build the dataset
samples = names(counts)
condition = factor(c(cond))
colData = data.frame(samples=samples, condition=condition)

# Create DESEq2 dataset.
dds = DESeqDataSetFromMatrix(countData=counts, colData=colData, design = ~condition)
#dds = DESeqDataSetFromMatrix(countData=counts, colData=colData, design = ~1)

#rlog transformation
#rld <- rlog(dds, blind=FALSE)

# Variance Stabilizing Transformation.
vsd = vst(dds)

# Open the drawing device.
pdf('pca.pdf', width = WIDTH, height = HEIGHT)

nudge = position_nudge(y = 1)
plotPCA(vsd, intgroup=c("condition")) + geom_label(aes(label = name), position = nudge)

dev.off()

library(pheatmap)
library("RColorBrewer")

sampleDists = dist(t(assay(vsd)))
sampleDistMatrix = as.matrix(sampleDists)

rownames(sampleDistMatrix) = vsd$samples
colnames(sampleDistMatrix) = NULL

colors = colorRampPalette( rev(brewer.pal(9, "Blues")) )(255)

# Open the drawing device.

pdf('heatmap.pdf', width = 8, height = HEIGHT)

pheatmap(sampleDistMatrix,
         clustering_distance_rows=sampleDists,
         clustering_distance_cols=sampleDists,
         col=colors)  + geom_label(aes(label = name))

dev.off()
```
