## Supplementary material for "Global Release of Translational Repression Across *Plasmodium’s* Host-to-Vector Transmission Event": Fig S1-6

Rios *et al.* (2024) Figure S1

A.

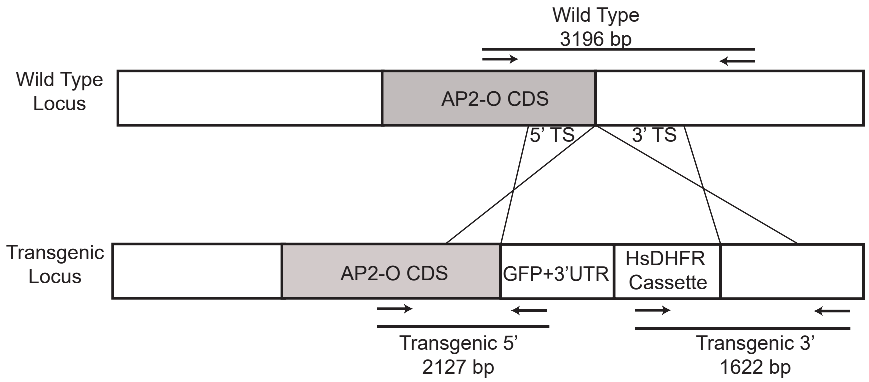


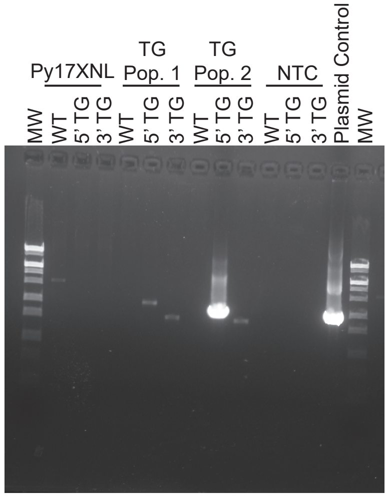


Py1534 - AP2-O::GFP

Rios *et al.* (2024) Figure S2

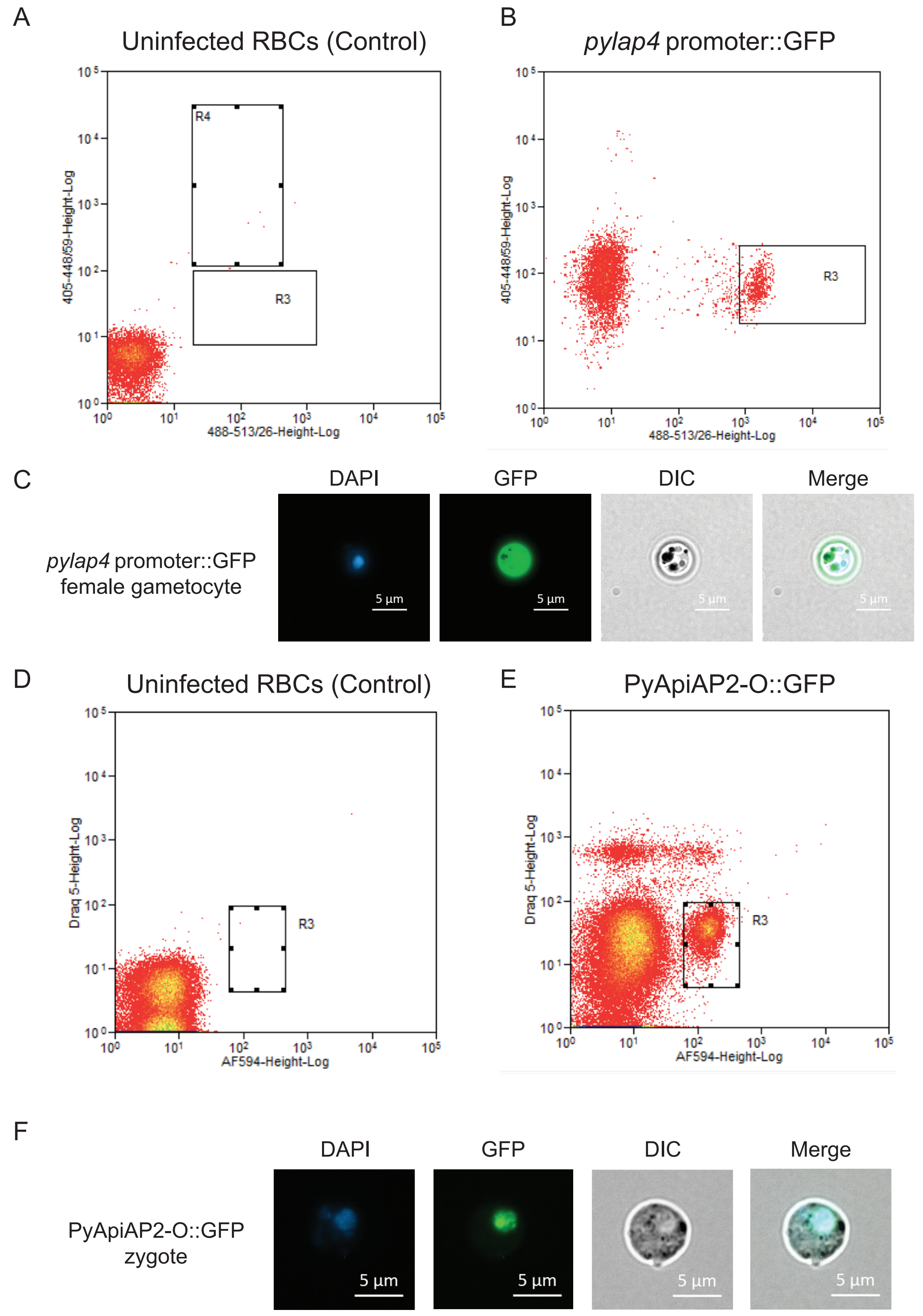

Rios *et al.* (2024) Figure S3

A

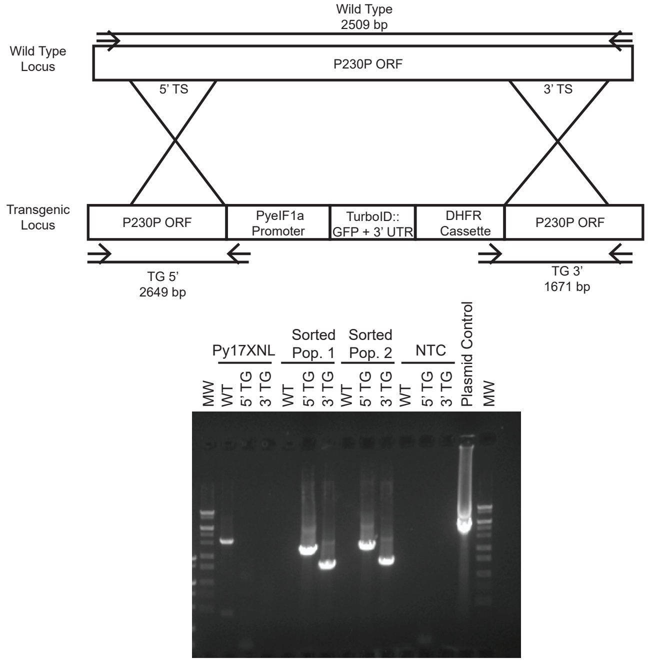


Py1419 unfused TurboID::GFP control

B

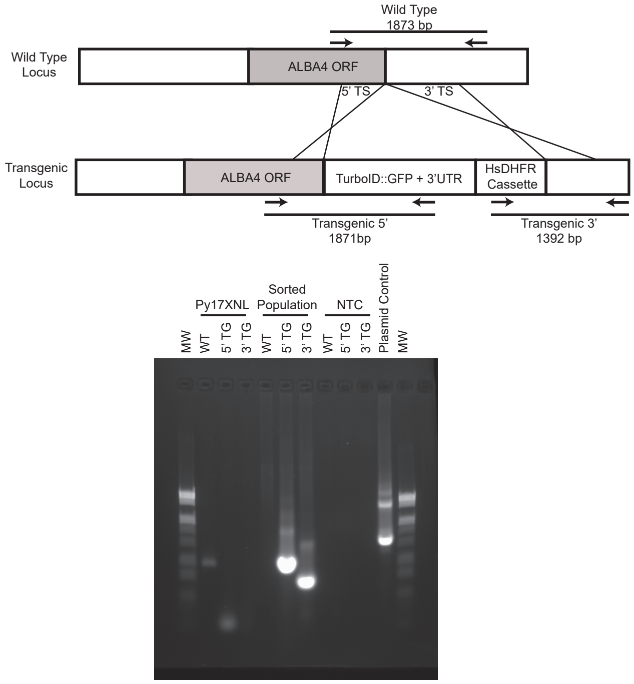


Py1385 ALBA4::TurboID::GFP

C

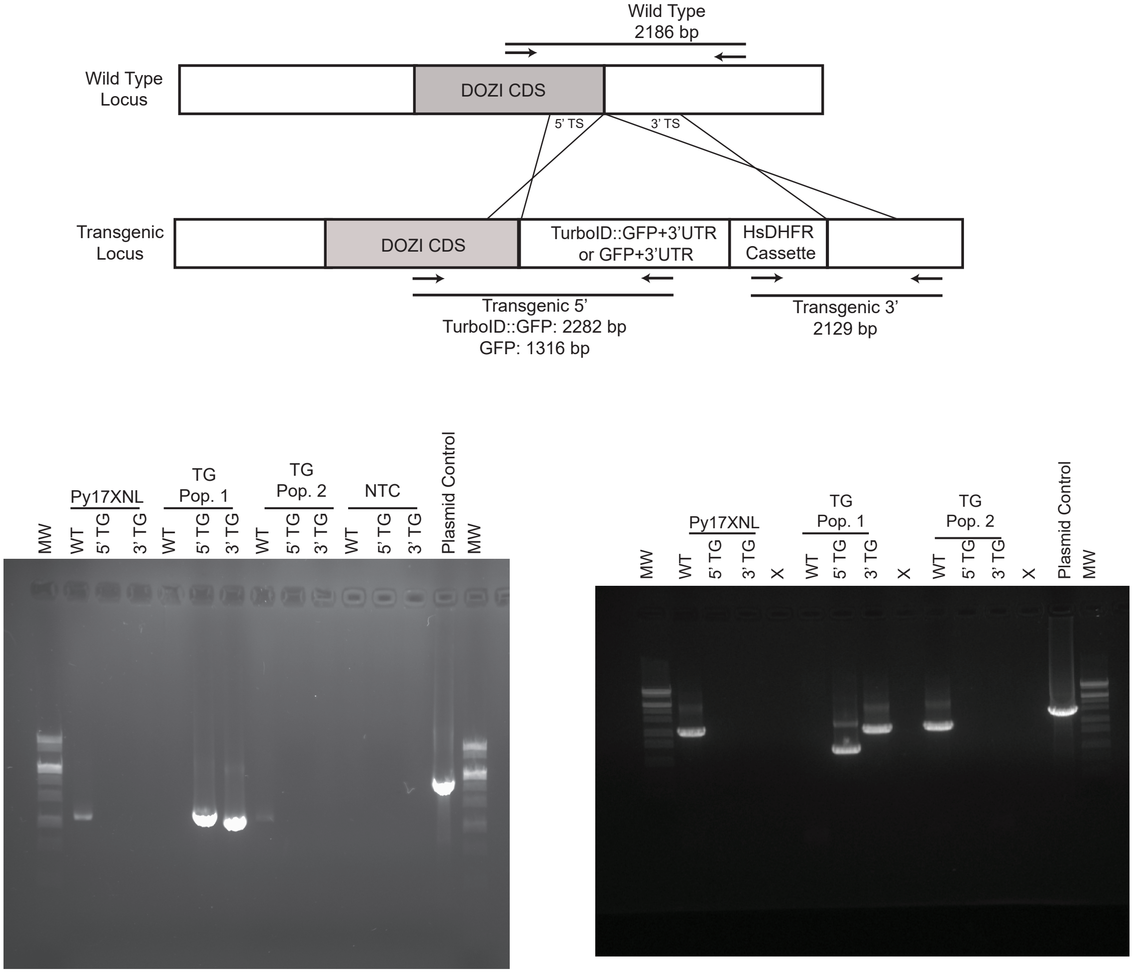


Py1490 DOZI::TurboID::GFP

Py1252 - DOZI::GFP

Rios *et al.* (2024) Figure S4

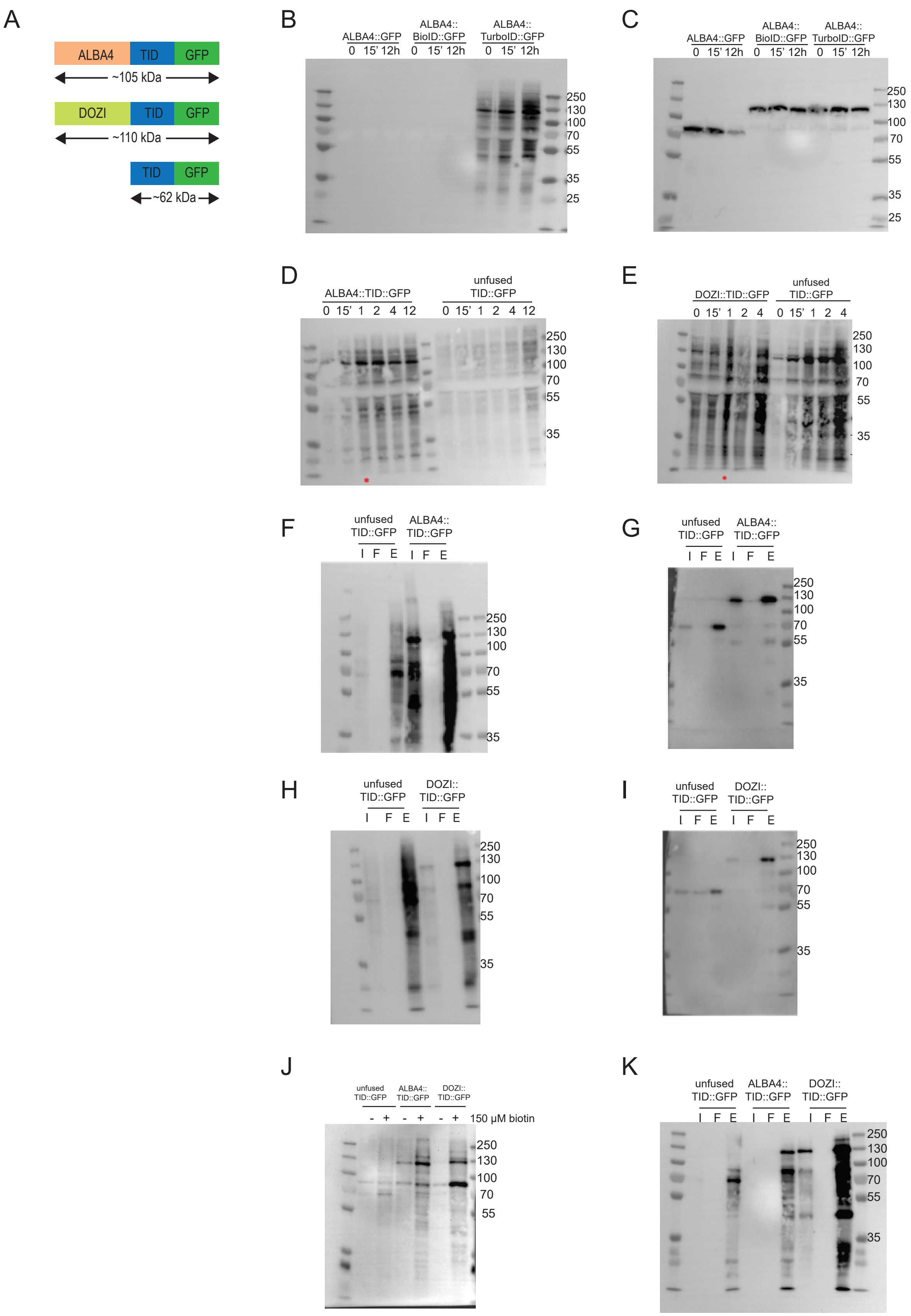

Rios *et al.* (2024) Figure S5

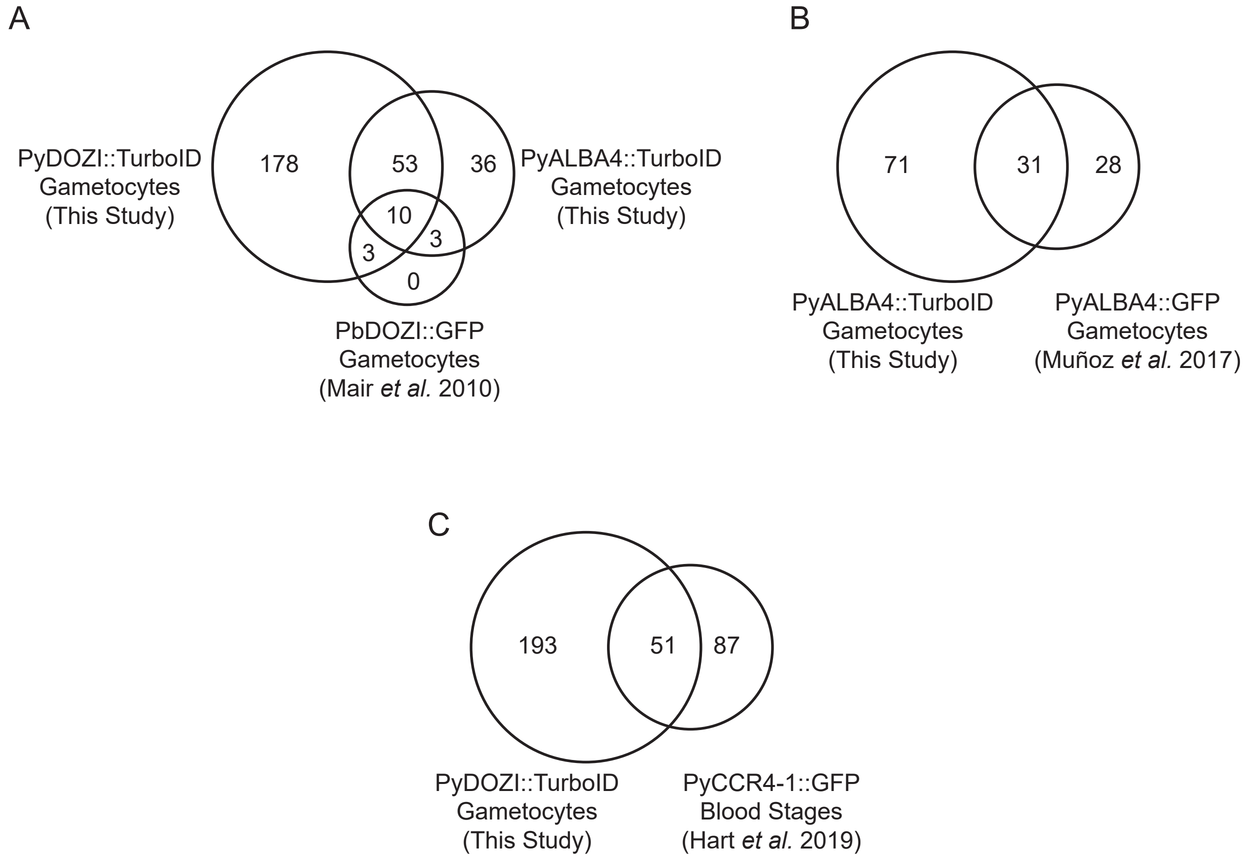

Rios *et al.* (2024) Figure S6

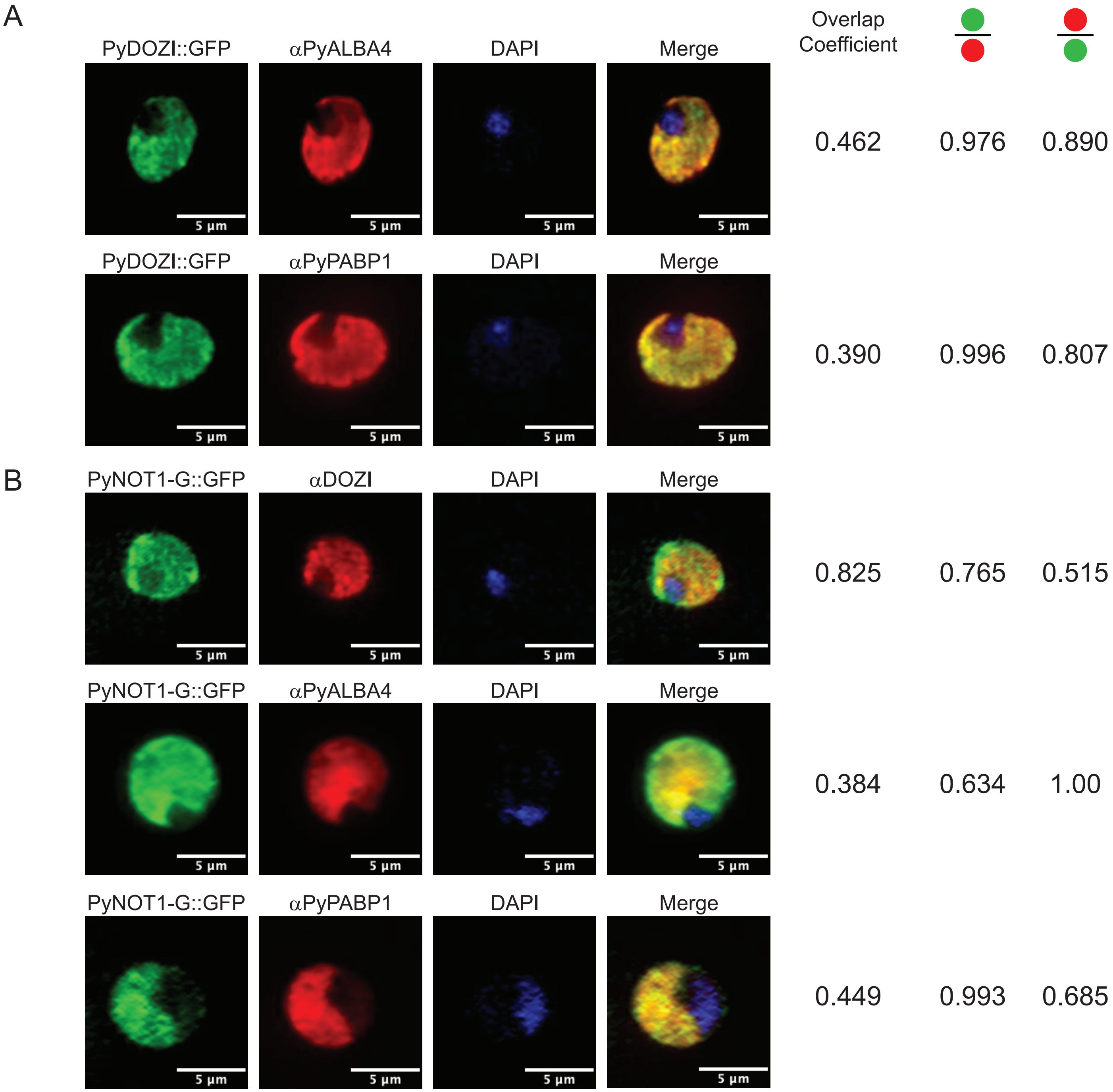
